# Adaptive preservation of orphan ribosomal proteins in chaperone-stirred condensates

**DOI:** 10.1101/2022.11.09.515856

**Authors:** Asif Ali, Rania Garde, Olivia C Schaffer, Jared A M Bard, Kabir Husain, Samantha Keyport Kik, Kathleen A Davis, Sofia Luengo-Woods, D Allan Drummond, Allison H Squires, David Pincus

**Affiliations:** Department of Molecular Genetics and Cell Biology, University of Chicago, Chicago, IL; Committee on Genetics, Genomics, and Systems Biology, University of Chicago, Chicago, IL; Pritzker School for Molecular Engineering, University of Chicago, Chicago, IL; Department of Biochemistry and Molecular Biology, University of Chicago, Chicago, IL; Department of Physics, University of Chicago, Chicago, IL; Department of Medicine, Section of Genetic Medicine, University of Chicago, Chicago, IL; Institute for Biophysical Dynamics, University of Chicago, Chicago, IL; Center for Physics of Evolving Systems, University of Chicago, Chicago, IL

## Abstract

Ribosome biogenesis is among the most resource-intensive cellular processes, with ribosomal proteins accounting for up to half of all newly synthesized proteins in eukaryotic cells. During stress, cells shut down ribosome biogenesis in part by halting rRNA synthesis, potentially leading to massive accumulation of aggregation-prone “orphan” ribosomal proteins (oRPs). Here we show that during heat shock in yeast and human cells, oRPs accumulate as reversible condensates at the nucleolar periphery recognized by the Hsp70 co-chaperone Sis1/DnaJB6. oRP condensates are liquid-like in cell-free lysate but solidify upon depletion of Sis1 or inhibition of Hsp70. When cells recover from heat shock, oRP condensates disperse in a Sis1-dependent manner, and their ribosomal protein constituents are incorporated into functional ribosomes in the cytosol, enabling cells to efficiently resume growth.

**One sentence summary:** During stress, molecular chaperones preserve “orphan” ribosomal proteins (RPs) – RPs that are not bound to rRNA – in liquid-like condensates, maintaining the RPs in a usable form and enabling cells to efficiently resume growth upon recovery from stress.

## INTRODUCTION

Cells must double their ribosome content each division cycle, and ribosome biogenesis may be rate-limiting for proliferation (Lempiainen and Shore, 2009; Maaløe and Kjeldgaard, 1966; Scott et al., 2014; Warner, 1999). In yeast, where eukaryotic ribosome biogenesis has been most thoroughly studied, ribosomal proteins (RPs) account for up to half of all newly synthesized proteins under nutrient-rich conditions (Ingolia et al., 2009; Shore and Albert, 2022). This huge investment of resources results in the synthesis of more than 10^5^ RPs each minute (Shore and Albert, 2022; Warner, 1999).

To assemble a mature yeast ribosome, 79 RPs must stoichiometrically associate with four ribosomal RNA (rRNA) molecules to form the large 60S and small 40S subunits (Woolford and Baserga, 2013). Many RPs cannot adopt their native structures and are highly aggregation prone in the absence of rRNA (Jakel et al., 2002). This poses a challenge in the compartmentalized eukaryotic cell: rRNA is transcribed in the nucleolus, while RPs are synthesized on ribosomes in the cytosol. Most RPs translocate from the cytosol into the nucleolus, and specific karyopherins and dedicated chaperones known as escortins mediate their transport (Pillet et al., 2017). Transcription of rRNA and the mRNAs encoding RPs is coordinately regulated and rapidly repressed upon nutrient limitation or other environmental stressors such as heat shock (Gasch et al., 2000; Gasch and Werner-Washburne, 2002; Sawarkar, 2022; Shore et al., 2021). However, while this transcriptional attenuation occurs even under modest stress conditions in yeast such as 37ºC, protein synthesis remains active and is only repressed at temperatures above 40ºC (Iserman et al., 2020; Muhlhofer et al., 2019). Given the rate of RP synthesis, such imperfect coordination between transcription and translation could result in a massive buildup of excess RPs relative to rRNA.

Several cellular mechanisms have been described that respond to “orphan” RPs (oRPs), i.e., RPs that are not bound to rRNA and/or otherwise mislocalized in the cell (Juszkiewicz and Hegde, 2018). Two distinct pathways, mediated by the Tom1/HUWE1 and UBE2O ubiquitin ligases, recognize oRPs and target them for proteasomal degradation (Sung et al., 2016; Yanagitani et al., 2017). Loss-of-function mutations in RP genes – which result in stoichiometric imbalance among RPs and accumulation of oRPs – lead to a class of human diseases known as “ribosomopathies” (Narla and Ebert, 2010). Mechanistically, the accumulation of oRPs in ribosomopathies is thought to result in the sequestration and inactivation of the ubiquitin ligase MDM2 which normally degrades p53, and the accumulation of p53 results in cell cycle arrest and apoptosis (James et al., 2014). In yeast, which lacks p53, genetic disruptions that result in the accumulation of oRPs activate a ribosome assembly stress response (RASTR) that results in downregulation of RP genes and induction of the heat shock response (HSR), a transcriptional regulon encoding chaperones and other protein homeostasis (proteostasis) factors (Albert et al., 2019; Pincus, 2020; Tye et al., 2019). While RASTR has not yet been linked to physiological or environmental stress, the ability of oRPs to activate the HSR suggests that ribosome assembly is under surveillance by the proteostasis network.

Here we demonstrate that oRPs accumulate and drive key early events in the cellular response to heat shock. We find that oRPs interact with the J-domain protein (JDP) Sis1, an essential regulator of the chaperone Hsp70. oRPs trigger the localization of Sis1 in yeast, and its homolog DnaJB6 in human cells, to the periphery of the nucleolus upon heat shock. Rather than targeting oRPs for degradation, Sis1 and Hsp70 maintain oRPs in dynamic condensates that remain liquid-like in cell-free lysate. Following recovery from heat shock, oRP condensates disperse, the RPs are incorporated into functional ribosomes in the cytosol, and cells efficiently resume proliferation. By actively maintaining condensate reversibility, Sis1 and Hsp70 preserve oRP functionality and conserve cellular resources during stress.

## RESULTS

### Sis1 localizes to the nucleolar periphery during heat shock

To define the molecular species that drive the spatial reorganization of the proteostasis network during heat shock, we focused on the J-domain protein (JDP) Sis1. Subcellular re-localization of Sis1 represents the earliest known cell biological event following heat shock in yeast (Feder et al., 2021). We constructed a four-color yeast strain to monitor Sis1 localization in live cells with respect to the nucleolus, the nuclear boundary, the cortical endoplasmic reticulum, and cytosolic protein aggregates marked by Hsp104 (Table S1). We imaged this strain in three dimensions over a heat shock time course using lattice light sheet microscopy (Video S1). Prior to heat shock, Sis1 was concentrated in the nucleoplasm and diffuse throughout the cytosol (Figure 1A). Acute heat shock at 39ºC resulted in re-localization of Sis1 from the nucleoplasm to a region surrounding the nucleolus within 2.5 minutes (Figure 1A). Image analysis revealed increased Sis1 peri-nucleolar localization in nearly all cells following 2.5 and 10 minutes of heat shock (Figure 1B; Figure S1A; see methods). We additionally observed overlapping cytosolic foci containing Sis1 and Hsp104 that formed with delayed kinetics compared to Sis1 peri-nucleolar localization (Figure 1A; Figure S1B). Ongoing protein synthesis and/or ribosome dynamics have been shown to contribute to activation of the heat shock transcriptional response and to re-localization of Sis1 and Hsp104 (Feder et al., 2021; Garde et al., 2022; Masser et al., 2019; Triandafillou et al., 2020; Tye and Churchman, 2021). Sis1 localization to the nucleolar periphery and cytosolic foci formation was abolished following heat shock upon pre-treatment with the translation elongation inhibitor cycloheximide (CHX) (Figure 1A,B; Figure S1B). Thus, ongoing translation drives Sis1 spatial dynamics during heat shock – in the cytosol and within the nucleus.

**Figure 1.**
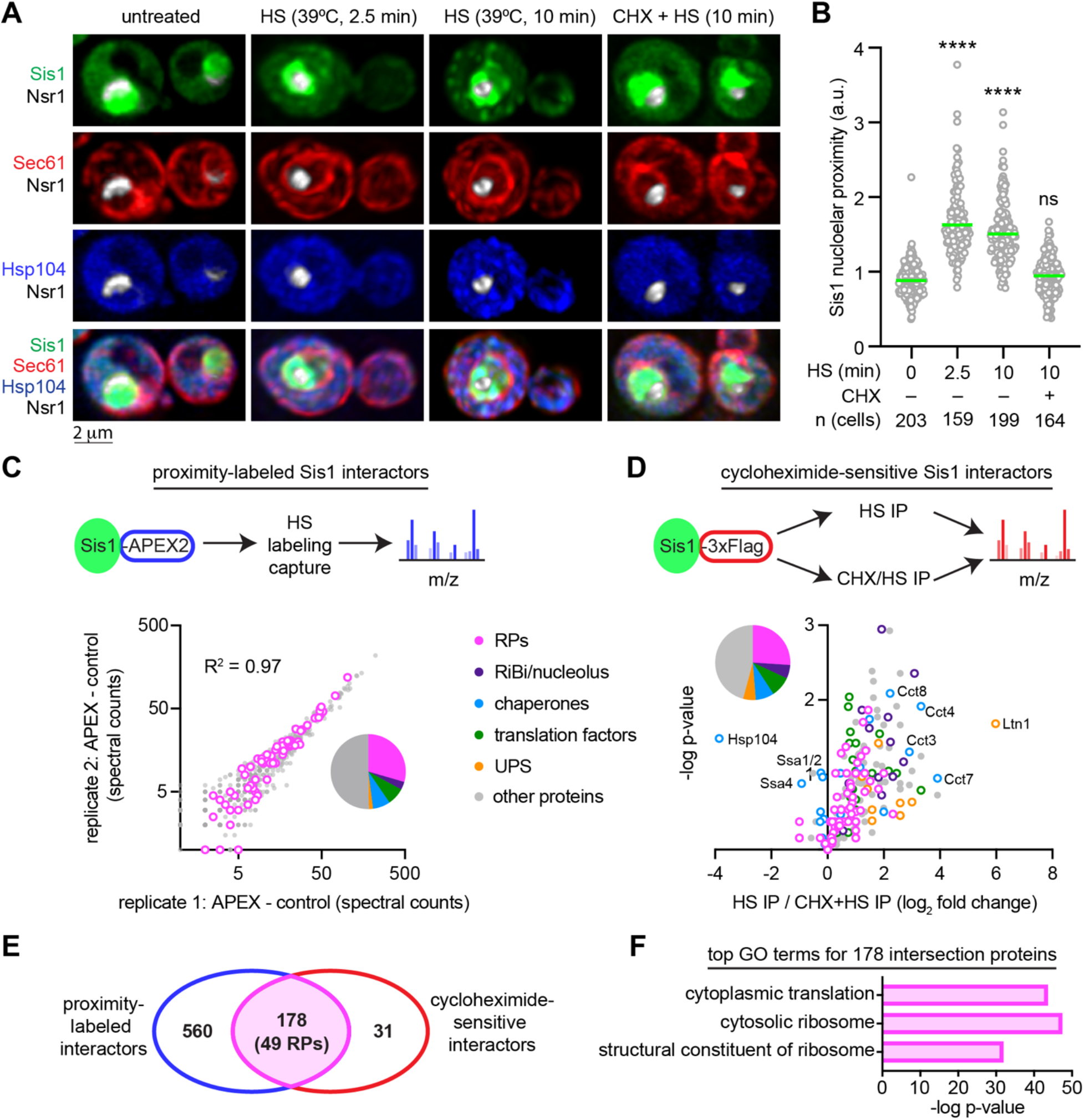
Sis1 localizes to the nucleolar periphery and interacts with ribosomal proteins during heat shock. **A)** Lattice light sheet live-imaging of yeast cells with endogenously tagged Sis1-mVenus (green), Hsp104-TFP (blue), Sec61-Halo (red) and Nsr1-mScarletI (white) under non-stress (30°C), heat shock (39°C, 2.5 and 10 min) and pre-treated with cycloheximide (CHX, 50μg/ml, 5min) followed by heat shock (39°C, 10 min). **B)** Single cell quantification of Sis1 nucleolar proximity defined as the ratio of mean Sis1 intensity in the half of the nucleus containing the nucleolus to the mean Sis1 intensity in the other half of the nucleus. Statistical significance was determined by Brown-Forsythe and Welch ANOVA test with multiple comparisons. \*\*\*\**P* < 0.0001, ns (non-significant). **C)** Top: Schematic of in vivo proximity labelling of Sis1-APEX2 followed by mass spectrometry analysis. Bottom: Enrichment of proteins labeled by Sis1-APEX2 following HS in two biological replicates. Ribosomal proteins highlighted in magenta. **D)** Top: Workflow to IP Sis1-3xFlag following heat shock (39°C, 10 min) pretreated with either vehicle or cycloheximide (50μg/ml, 5 min). Bottom: Volcano plot demonstrating magnitude and statistical significance of cycloheximide sensitivity of Sis1 interactors. **E)** Venn diagram of proteins identified in (C) and (D). **F)** Gene ontology terms enriched among the 178 intersection proteins from (E).

### Ribosomal proteins are a major class of Sis1 interactors during heat shock

To identify proteins that Sis1 interacts with during heat shock, we utilized proximity-dependent labeling and affinity capture coupled to mass spectrometry (MS). To achieve this, we endogenously fused Sis1 to the engineered peroxidase APEX2 (Hung et al., 2016). We heat shocked cells expressing Sis1-APEX2 and performed the proximity labeling in the presence and absence of a pulse of H_2_O_2_ to distinguish specifically and nonspecifically labeled proteins (see methods). Biological replicates showed strong concordance in the abundance of the 698 proximity labeled proteins reproduced across replicates and quantifiable above background (Figure 1C; Table S2: APEX data). The data revealed that during heat shock, Sis1 is proximal to other chaperones, metabolic enzymes, translation factors, ribosome biogenesis factors, and – most prominently – ribosomal proteins (RPs) (Figure 1C; Figure S1C).

To complement the proximity ligation approach, we performed immunoprecipitation (IP)-MS of Sis1-3xFlag following heat shock in the absence and presence of CHX. Of the 227 proteins we reproducibly pulled down during heat shock with Sis1, 209 were depleted in the CHX-treated cells relative to cells that were heat shocked in the absence of CHX (Figure 1D; Figure S1D; Table S3: IP-MS data). Notably, interactions between Sis1 and cytosolic chaperones such as Hsp70 and Hsp104 increased in cells treated with CHX, while interactions with proteostasis factors involved in co-translational protein folding and degradation – including members of the TRiC/CCT chaperonin complex and the ubiquitin ligase Ltn1 – were sensitive to CHX. However, the most abundant class of CHX-sensitive Sis1 interactors was RPs. Comparison of the set of APEX2-labeled Sis1 interactors with the set of CHX-sensitive Sis1 interactors revealed an intersection of 178 proteins, 49 of which are RPs (Figure 1E,F). These proteomic results indicate that RPs are a major class of CHX-sensitive Sis1 interactors.

### Sis1 interacts with orphan RPs at the nucleolar periphery upon heat shock

The CHX-sensitivity of the interaction between Sis1 and RPs could result either from interactions of Sis1 with mature ribosomes or from interactions of Sis1 with newly synthesized RPs. To distinguish these possibilities, we performed pulse-chase experiments to selectively label either mature or new RPs. We endogenously tagged RPs of the large (Rpl26a, Rpl25) and small ribosomal subunits (Rps4b, Rps9a) with the HaloTag to enable labeling with cell-permeable haloalkanes, either a non-fluorescent “blocker” (7-bromoheptanol) or fluorescent dye (JF646) (Figure 2A). To label mature RPs, we first pulsed with JF646 and subsequently chased with blocker to mask any new RPs. Conversely, we first added blocker to mask all the mature RPs and then added JF646 to label new RPs. We pulled down Sis1-3xFlag under these two labeling schemes in the presence or absence of heat shock. Despite the great excess of mature RPs in the input samples relative to the new RPs, Sis1 co-precipitated only with the new RPs and only during heat shock (Figure 2B,C; Figure S2A).

**Figure 2.**
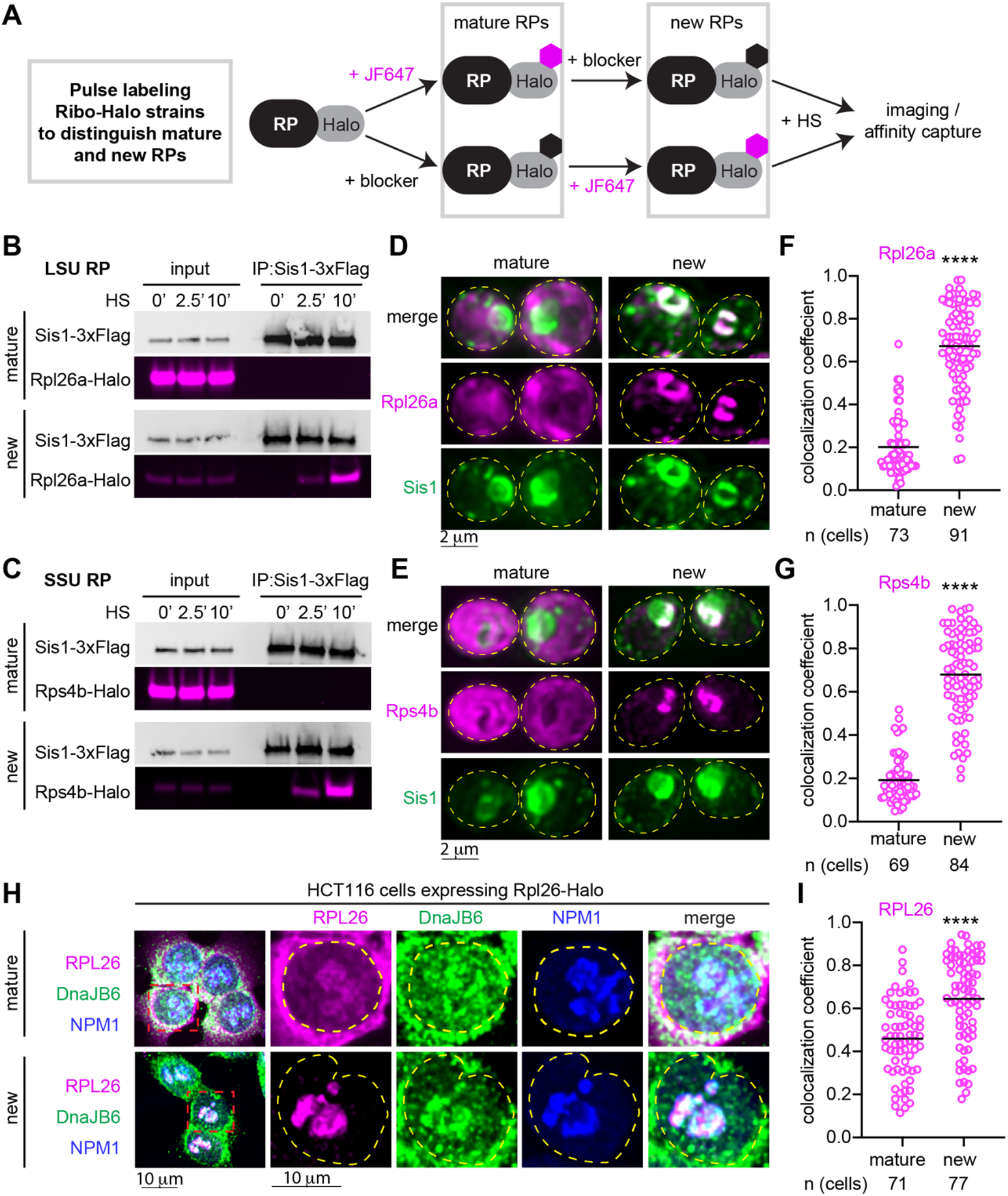
Orphan ribosomal proteins interact with Sis1/DnaJB6 at the nucleolar periphery. **A)** Workflow for in vivo pulse-labelling of mature and new ribosomal proteins. **B)** IP of Sis1-3xFlag and mature or new Rpl26a-Halo from cells heat shocked at 39°C for the indicated times. **C)** As in (B), but for Rps4b-Halo. **D)** Lattice light sheet live imaging of yeast under heat shock (39°C, 10 min) expressing Sis1-mVenus and labeled for either new or mature Rpl26a-Halo. The dashed line indicates the cellular boundary. **E)** As in (D), but for Rps4b-Halo. **F)** Colocalization (Mander’s overlap coefficient) of Sis1-mVenus with either mature or new Rpl26a-Halo in heat shocked cells (39°C, 10 min). **G)** As in (F), but for Rps4b-Halo. *P* values were calculated with a two-tailed Welch’s t-test. \*\*\*\**P* < 0.0001. **H)** Human HCT116 cells stably expressing RPL26-Halo labelled for mature or new RPL26 and heat shocked (43°C for 30 min). Cells were fixed and immunostained for DnaJB6 and NPM1. Dashed line indicates the nuclear boundary. **I)** Colocalization (Mander’s overlap coefficient) of DnaJB6 with either mature or new RPL26-Halo in heat shocked cells (43°C, 30 min). *P* values were calculated with a two-tailed Welch’s t-test. \*\*\*\**P* < 0.0001.

Next, we imaged RPs with respect to Sis1. We endogenously tagged three large subunit RPs (Rpl26a, Rpl25, and Rpl29) and three small subunit RPs (Rps4b, Rps9a, and Rps3) with the HaloTag in the Sis1-mVenus background. Using the pulse-chase labeling scheme described above, we imaged either mature or new RPs in live cells. Under nonstress conditions, all newly synthesized RPs were immediately localized to the cytosol, demonstrating the rapidity of ribosome biogenesis (Figure S2B). We heat shocked the cells for 10 minutes to cause both Sis1 cytosolic foci formation and peri-nucleolar Sis1 accumulation. Mature RPs formed neither cytosolic foci nor peri-nucleolar structures, and the Sis1 cytosolic foci localized away from mature ribosomes (Figure 2D-G; Figure S2C-F). By contrast, new Rpl26a, Rpl25, Rps4b and Rps9b – which are all incorporated into ribosomal subunits in the nucleolus (Woolford and Baserga, 2013) – accumulated at the nucleolar periphery as mis-localized oRPs that colocalized with Sis1 (Figure 2D-G; Figure S2C,D). However, new Rpl29 and Rps3 – which are incorporated into their respective ribosomal subunits at a late stage of biogenesis which occurs in the cytosol (Woolford and Baserga, 2013) – neither localized to the nucleolar periphery nor co-localized with Sis1 (Figure S2E,F). These observations suggest that oRPs accumulate at the nucleolar periphery with Sis1 during heat shock provided that the RP is incorporated into ribosome assembly in the nucleolus.

### DnaJB6 co-localizes with newly synthesized RPL26 in human cells during heat shock

Do oRPs also accumulate during heat shock in human cells? To test this, we obtained HCT116 human cells with a Halo-tagged copy of RPL26 integrated into the genome (An et al., 2020). The human JDP DnaJB6 is likely to be the functional homolog of Sis1 in human cells (Klaips et al., 2020). To determine whether DnaJB6 co-localizes with RPs during heat shock, we followed a similar pulse-labeling protocol in the HCT116 cells as we used in yeast (Figure 2A), fixed the cells following heat shock, and stained with antibodies against DnaJB6 and NPM1 to mark the periphery of the nucleolus. As in yeast, mature RPL26 was primarily localized to the cytosol, though some was detectible colocalizing with NPM1 (Figure 2H). Mature RPL26 showed partial colocalization with DnaJB6 during heat shock (Figure 2H,I). However, as in yeast, newly synthesized RPL26 was highly concentrated at the periphery of the nucleolus and colocalized with DnaJB6 (Figure 2H,I). Thus, in yeast and human cells, heat shock triggers the peri-nucleolar accumulation of oRPs that colocalize with homologous JDPs.

### RP production is required for peri-nucleolar recruitment of Sis1/DnaJB6

While we have implicated the accumulation of oRPs in recruiting Sis1 to the nucleolar periphery during heat shock, we have not yet established whether oRPs are necessary for Sis1 re-localization. To test this, we utilized auxin inducible degradation to acutely deplete Ifh1, a transcription factor required for the expression of RPs in yeast (Schawalder et al., 2004). Depletion of Ifh1 impaired cell growth, and mRNA deep sequencing under nonstress conditions revealed that acute loss of Ifh1 resulted in reduced expression of RP mRNAs with few other changes to the transcriptome (Figure 3A,B). Upon heat shock, cells depleted for Ifh1 showed reduced transcriptional induction of Hsf1-regulated genes without alterations in other stress response genes (Figure 3C; Figure S3A,B). On average, Hsf1 targets were reduced by 25% in Ifh1-depleted cells, suggesting that newly synthesized RPs contribute to activation of the heat shock response. While the effect of Ifh1-depletion on the heat shock response was relatively modest, loss of Ifh1 nearly abolished Sis1 localization to the nucleolar periphery during heat shock in most cells (Figure 3D,E). However, Ifh1 depletion did not prevent formation of Sis1 cytosolic foci during heat shock, demonstrating a specific effect on peri-nucleolar Sis1 (Figure S3C,D). In HCT116 cells, we used the mTOR inhibitor torin-1 to reduce expression of RPs (Thoreen et al., 2012), and this treatment prevented heat shock induced localization of DnaJB6 to the nucleolar periphery in most cells (Figure 3F,G). These data suggest that oRPs recruit Sis1/DnaJB6 to the nucleolar periphery during heat shock.

**Figure 3.**
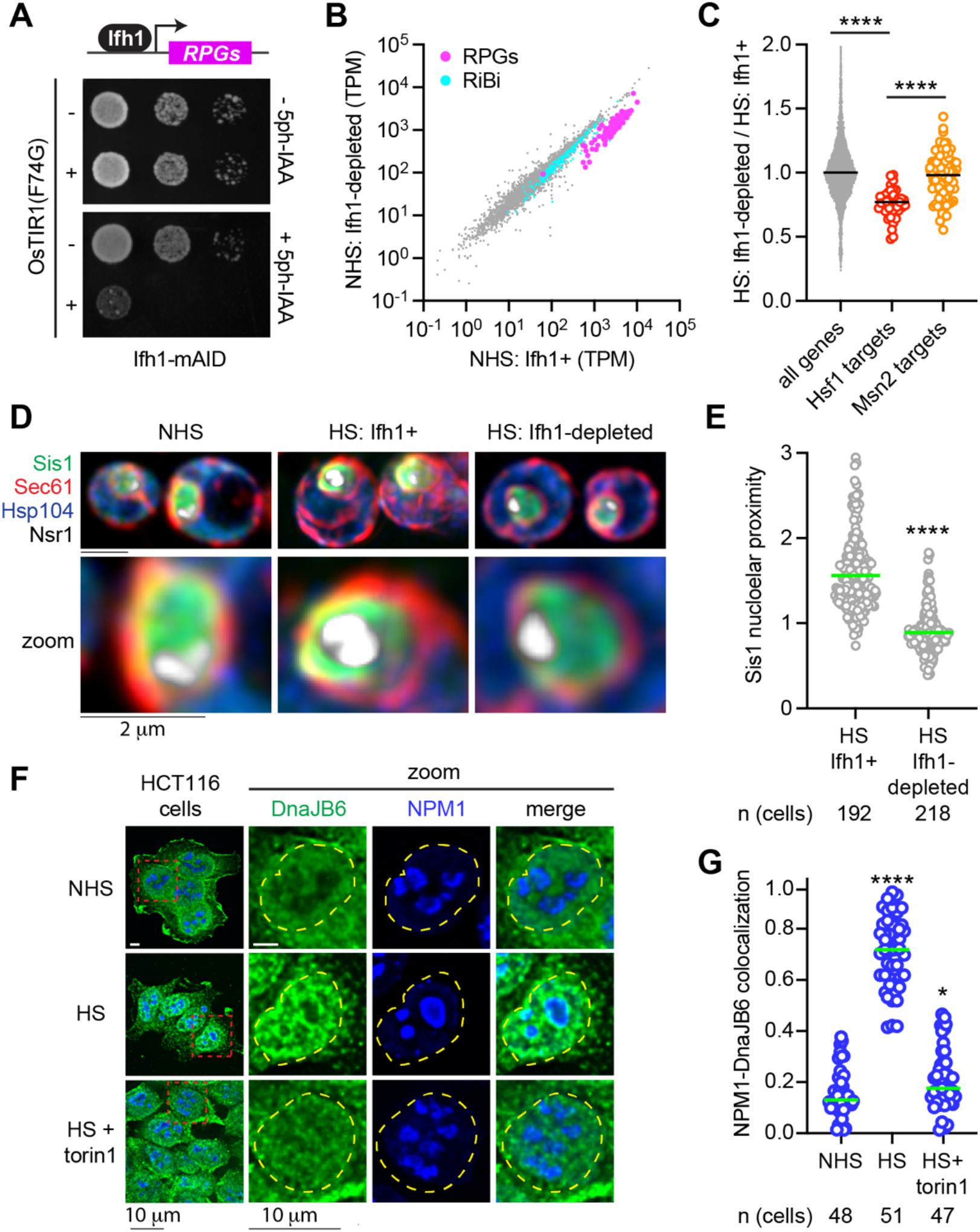
Ribosomal proteins drive Sis1/DnaJB6 localization to the nucleolar periphery. **A)** Dilution series spot assay of yeast cells expressing Ifh1-mAID with or without estradiol inducible OsTIR1(F74G) spotted on to rich medium supplemented with either estradiol (1μM) alone or with 5ph-IAA (5μM) grown for 48 hr. **B)** Scatter plot of RNA seq data from Ifh1-mAID/OsTIR1(F74G) treated with estradiol alone or along with 5phIAA for 30 min. Ribosome biogenesis factors (RiBi, cyan) and ribosomal protein genes (RPGs, magenta) are highlighted. NHS (30°C) **C)** Ifh1-dependent fold change in gene expression in heat shocked cells (39°C, 5 min) showing a reduction in Hsf1 target gene induction (red dots) but not in Msn2 target gene induction (yellow dots). **D)** LLS imaging of the spatial distribution of Sis1-mVenus in non-stressed cells (NHS, 30°C), heat shocked cells (HS, 39°C, 5 min), and heat shocked cells depleted for Ifh1. **E)** Quantification of Sis1 nucleolar proximity during heat shock in cells with Ifh1 and following Ifh1 depletion. Statistical significance was determined by Brown-Forsythe and Welch ANOVA test with multiple comparisons. \*\*\*\**P* < 0.0001, ns (non-significant). **F)** Immunostaining of HCT116 cell lines for DnaJB6 (green) and NPM1 (Blue) under nonstressed, heat shocked (43°C, 30 min) and heat shock of cells pretreated with Torin1 (300nM, 30 min). **G)** Quantification of colocalization (Mander’s overlap coefficient) of NPM1 and DnaJB6 in single cells in the conditions in (F). Statistical significance was determined by Brown-Forsythe and Welch ANOVA test with multiple comparisons. \*\*\*\**P* < 0.0001, \**P* =0.0248

### oRPs form dynamic condensates with Sis1 during heat shock

We next performed time-lapse 3D lattice light sheet imaging of newly synthesized Rpl26a to observe spatiotemporal dynamics of oRPs during heat shock with respect to the nucleolus and Sis1. Over a sustained 10-minute heat shock, new Rpl26a and Sis1 remained localized at the nucleolar periphery in dynamic clusters that rapidly reorganized via fission and fusion (Video S2; Figure 4A). The fluorescence signal remained nearly constant over time, suggesting that these clusters are stable and are not sites of degradation (Figure S4A,B). We will refer to these clusters as “oRP condensates”. To probe the sensitivity of oRP condensates to alcohols known to disrupt certain other biomolecular condensates, we treated cells with 2,5-hexanediol (HD) and 1,6-HD at carefully optimized concentrations (Chowdhary et al., 2022). 2,5-HD had no effect on Sis1 or new Rpl26a localization (Figure 4B). However, while treatment with 1,6-HD did not disrupt Rpl26a clustering, it resulted in loss of Sis1 peri-nucleolar localization (Figure 4B). This differential sensitivity to 1,6-HD suggests that the interactions that target Sis1 to oRPs are biophysically distinct from the interactions among the oRPs themselves. Remarkably, Sis1 localization to oRPs was restored upon washout of 1,6-HD (Figure 4B).

**Figure 4.**
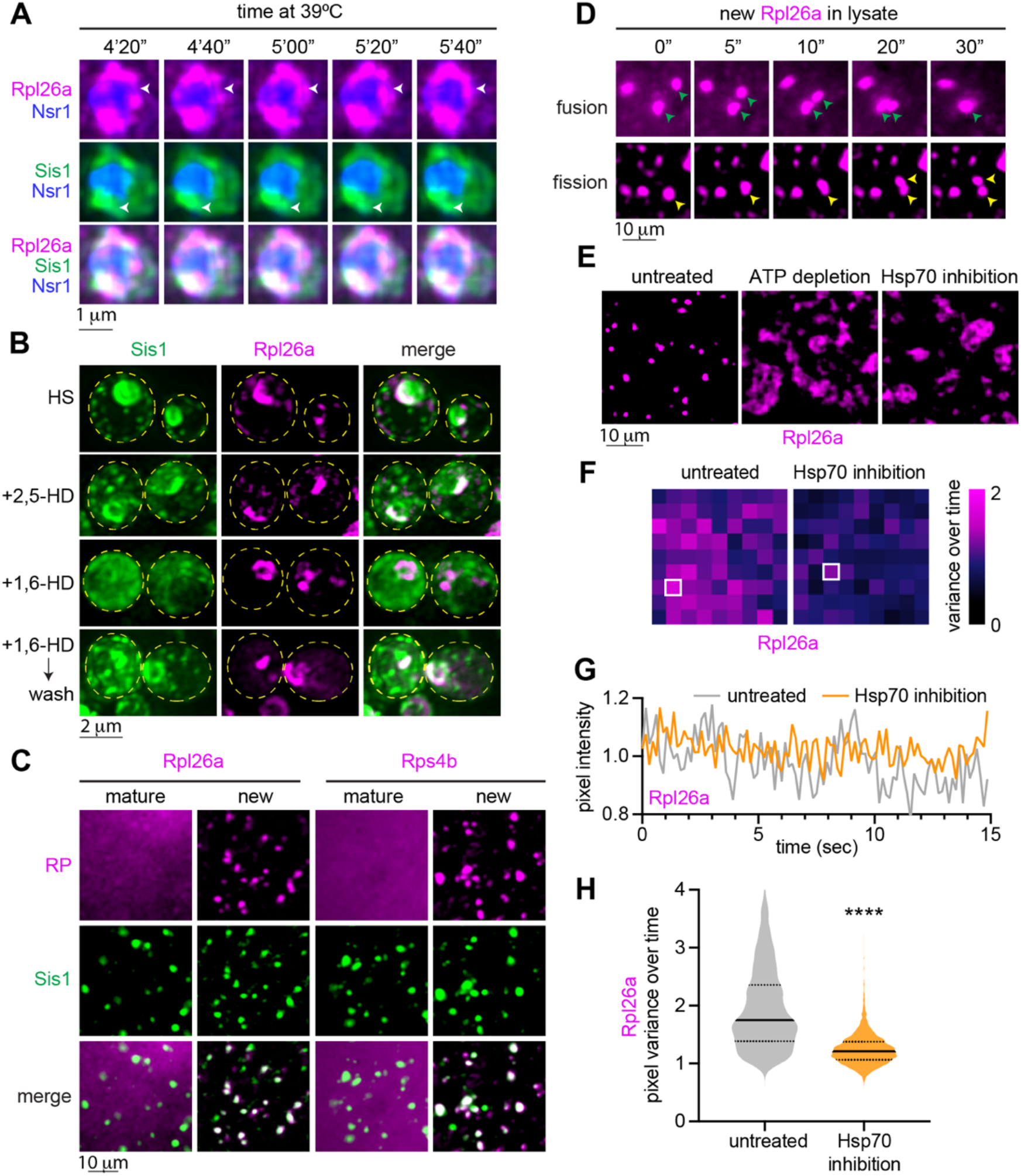
oRPs form dynamic condensates that are stable in cell-free extracts. **A)** 4D LLS imaging of Sis1-mVenus (green) and oRpl26a (magenta). **B)** Effect of 1,6-HD and 2,5-HD on Sis1 localization and oRP condensates. Cells were heat shocked (39°C) for 10 min and incubated with either 5% 1,6 HD or 2,5 HD for 2 min. 1,6-HD was washed out for an additional 2 min at heat shock temperature and cells were imaged again (last panel). **C)** Imaging of mature and new Rpl26a-Halo and Rps4b-Halo along with Sis1-mVenus in cell-free lysate from heat shocked cells (39°C, 10min). **D)** Time lapse imaging of heat shocked cell-free lysate depicting fission and fusion of oRpl26a droplets. **E)** Depletion of ATP (apyrase, 0.05units/μl) or inhibition of Hsp70 (VER155008, 50μM) activity in the lysate resulted in the formation of irregular clumps of oRpl26a. **F)** Heat map representing the normalized variance of pixel photon count over time within the heat shock induced oRpl26a condensate and upon Hsp70 inhibition. **G)** Shot noise-normalized intensity over time of the most variable pixel from an oRpl26a condensate in lysate with vs. without Hsp70 inhibition. **H)** Distribution of normalized pixel variance of oRpl26a in condensates over time in lysate with and without Hsp70 inhibition. *P* values were calculated with a two-tailed Welch’s t-test. \*\*\*\**P* < 0.0001.

### Hsp70 activity maintains the liquid-like dynamics of oRP condensates in cell-free extract

To interrogate the biochemical and biophysical properties of oRP condensates, we developed an assay to observe them in crude cell lysate. We labeled mature and new Rpl26a and Rps4b in cells co-expressing Sis1-mVenus, heat shocked the cells, and resuspended cryo-milled lysate in buffer with a chemical composition similar to the yeast cytoplasm (see methods). In the lysate, mature RPs showed diffuse signal, but new Rpl26a and Rps4b were concentrated in round condensates which colocalized with Sis1-mVenus (Figure 4C). Condensates were nearly absent in lysate from non-heat shocked cells (Figure S4B). Incubation with RNA-specific dye revealed that the condensates appear to exclude RNA (Figure S4C). Moreover, addition of RNAse to the lysate did not disrupt condensate morphology or number, suggesting that the RPs are orphaned from rRNA (Figure S4D,E). We observed both fusion and fission of oRP condensates in the lysate (Figure 4D). To determine whether maintenance of oRP condensates required ATP, we added apyrase to hydrolyze ATP in the lysate. ATP depletion resulted in loss of round condensates and the formation of larger amorphous aggregates (Figure 4E). Since Sis1 activates ATP hydrolysis by Hsp70, we wondered whether Hsp70 activity might contribute to the requirement for ATP. Indeed, addition of the Hsp70 inhibitor VER-155008 phenocopied ATP depletion, resulting in the formation of large irregular clumps of Rpl26a (Figure 4E).

We collected videos of the oRP condensates in lysate in the presence and absence of the Hsp70 inhibitor to evaluate their homogeneity and measure the dynamics of oRPs moving within the condensates. Each pixel recording a sample with a time-invariant brightness distribution, such as a solid sample or a homogeneous liquid, is expected to exhibit a variance in photon count equal to its average photon count due to the underlying Poisson statistics of arriving photons (Lakowicz, 2006). Samples that are not time-invariant, such as an inhomogeneous liquid, will exhibit higher-than-expected variance at each pixel. We quantified the Poisson-normalized variance in Rpl26a fluorescence over time for each pixel as a metric indicating the movement of non-homogeneous material within a condensate (see methods: Condensate Pixel Variance Analysis) (Linsenmeier et al., 2022). High normalized variance (>1) indicates that a condensate is dynamically rearranging and therefore more liquid-like, while normalized variance close to 1 indicates that a condensate is less dynamic and more solid-like. Pixels in individual condensates showed large variance in untreated lysate and much lower variance in the Hsp70-inhibited lysate (Figure 4F,G). Collective analysis across all pixels revealed that Hsp70 inhibition significantly reduced the pixel variance (Figure 4H). Thus, Hsp70 inhibition in lysate makes oRP condensates less dynamic and more solid-like.

### oRP condensates are reversible upon recovery from heat shock

The liquid-like character of the oRP condensates in lysate prompted us to ask whether they might be reversible upon recovery from stress in cells. We pulse labeled new Rpl26a and Rps4b with JF646, heat shocked the cells for 5 minutes, and chased with excess blocker in media either at 39ºC for a sustained heat shock or 30ºC to monitor recovery (Figure 5A). Imaging revealed that while the oRP condensates persisted at 39ºC, cells allowed to recover at 30ºC showed Rpl26a and Rps4b signal disperse to the cytosol within 7.5 minutes (Figure 5B-D; Figure S5A,B). The fluorescence signal of Rpl26a and Rps4b localized at the nucleolar periphery during heat shock can be quantitatively accounted for in the cytosol after recovery, suggesting there is little to no degradation of oRPs (Figure S5C). We observed the same re-localization of the signal of pulse-labeled RPL26 from the nucleolar periphery to the cytosol in human HCT116 cells following recovery (Figure 5G,H).

**Figure 5.**
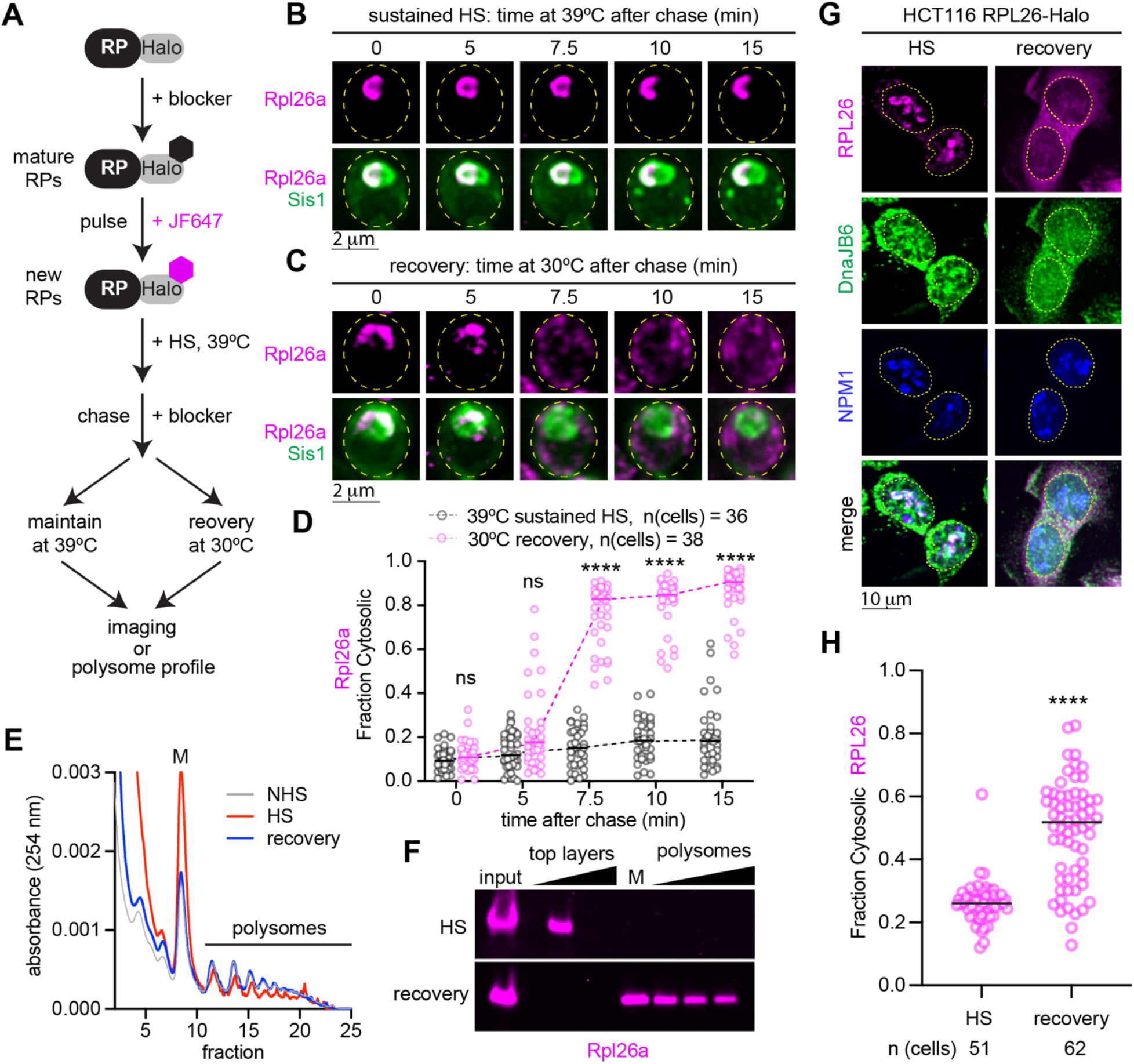
oRP condensates are reversible upon recovery from heat shock. **A)** Workflow to evaluate the fate of oRPs during sustained heat shock or recovery. **B)** Representative live-cell time lapse images of the spatial distribution of oRpl26a (magenta) and Sis1-mVenus (green) during sustained heat shock. **C)** As in (B) but following recovery from heat shock. **D)** Quantification of the fraction of cytosolic Rpl26a signal under sustained HS or recovery. Statistical significance was determined by Brown-Forsythe and Welch ANOVA test with multiple comparisons. \*\*\*\**P* < 0.0001, ns (non-significant). **E)** Polysome profiles of yeast expressing Rpl26a-Halo during nonstress, heat shock and recovery conditions. **F)** In-gel fluorescence of oRpl26a across the polysome profile in heat shock (39°C, 10 min) and recovery (30°C, 30 min). **G)** Localization of oRPL26 (magenta) in HCT116 cells following heat shock (43°C, 15 min) and recovery (37°C, 1 hr). Cells were fixed and immunostained for DnaJB6 (green) and NPM1 (blue). **H)** Quantification of the fraction of cytosolic RPL26 in HCT116 cells under sustained HS or recovery as in (G).

Given the cytosolic localization of pulse labeled RPs following recovery, we hypothesized that the RPs were being incorporated into mature ribosomes. To test this, we performed polysome profiling of lysate from unstressed yeast cells, heat shocked cells, and cells that been heat shocked and allowed to recover. Consistent with previous reports (Cherkasov et al., 2013; Grousl et al., 2013), we observed an increase in monosomes and a decrease in polysomes in heat shocked cells compared to unstressed cells that was completely reversed upon recovery (Figure 5E). In the heat shocked sample, we observed pulse labeled Rpl26a in the top layer fraction of cellular components not incorporated in or associated with ribosomes, consistent with it being an oRP (Figure 5F). By contrast, Rpl26a sedimented in the heavy polysome fractions in cells allowed to recover after heat shock (Figure 5F). Together, the cytosolic localization and co-sedimentation with polysomes suggest that RPs orphaned at the nucleolar periphery during heat shock are incorporated into functional ribosomes upon recovery.

### oRP condensate reversibility requires Sis1 and promotes fitness

Since Sis1 localizes to the oRP condensates, and Hsp70 activity was required to maintain the liquid-like character of the oRP condensates in lysate, we next asked whether the Sis1/Hsp70 chaperone system was also required for the dispersal of the condensates when cells recover from heat shock. Rather than mutating or inhibiting Hsp70 – which would have pleiotropic effects due to the diversity of processes in which Hsp70 participates – we utilized the “anchor away” approach we had previously established to conditionally deplete Sis1 from the nucleus (Feder et al., 2021). We generated RP-Halo strains in the Sis1 anchor away background and either left the cells untreated (Sis1+) or depleted Sis1 from the nucleus (Sis1-depleted). Then we pulse labeled new Rpl26a and Rps4b, heat shocked the cells at 39ºC, returned the cells to the recovery temperature of 30ºC, and imaged the RPs with respect to the nucleolus over time. In the Sis1+ condition, Rpl26a and Rps4b rapidly dispersed from the nucleolar periphery into the cytosol (Figure 6A,B; Figure S6A,B). However, in the Sis1-depleted cells, the RPs failed to disperse to the cytosol and remained adjacent to the nucleolus upon recovery (Figure 6A,B; Figure S6A,B). Consistent with a connection between the reversibility of the oRP condensate and their liquid-like internal dynamics, analysis of pixel-by-pixel normalized variance showed that new Rpl26a formed amorphous solid-like aggregates in lysate from Sis1-depleted cells (Figure 6C,D). These data suggest that Sis1 recruitment to the nucleolar periphery during heat shock contributes to maintaining the liquid-like properties of oRP condensates, which in turn enable them to be rapidly dispersed and utilized to restart ribosome biogenesis upon recovery from stress.

**Figure 6.**
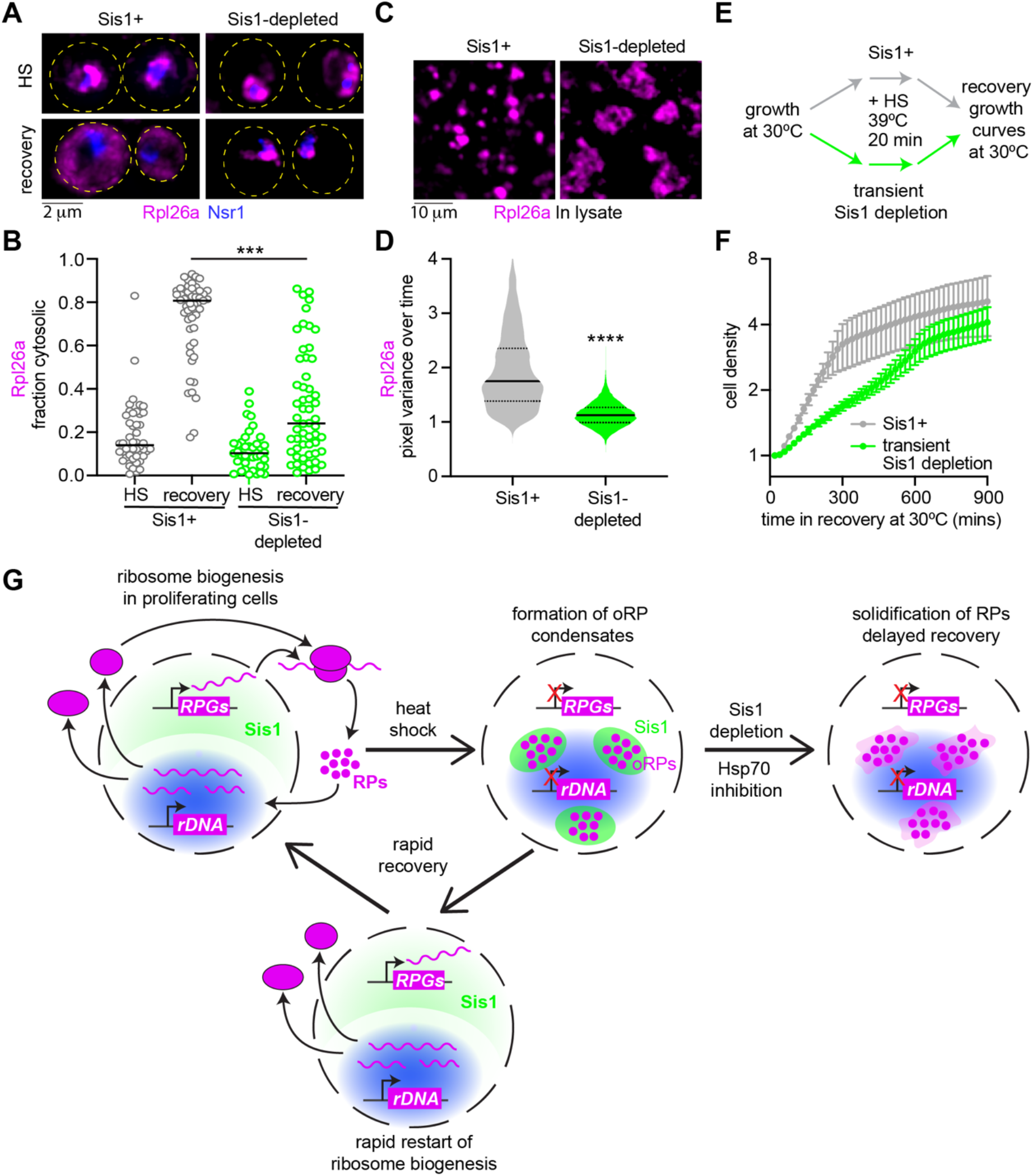
Sis1 promotes oRP condensate reversibility and growth following recovery. **A)** LLS imaging of oRPl26a (magenta) and the nucleolar marker Nsr1 (blue) during heat shock and recovery in the absence or presence of Sis1 depletion (10 min prior to heat shock). **B)** Quantification of the fraction of cytosolic Rpl26a under sustained HS or recovery in the absence or presence of Sis1 depletion. *P* values were calculated with a two-tailed Welch’s t-test. \*\*\*\**P* < 0.0001. **C)** Depletion of Sis1 resulted in formation of irregular clumps of Rpl26a in lysate. **D)** Distribution of normalized pixel variance of Rpl26a in condensates over time in control lysate and lysate with Sis1 depletion. Control lysate distribution replotted from Figure 4A. *P* values were calculated with a two-tailed Welch’s t-test. \*\*\*\**P* < 0.0001. **E)** Workflow of growth assay following recovery. **F)** Growth curve of cells following recovery from heat shock in the absence or presence of transient Sis1 depletion. Mean and standard deviation of 3 biological replicates are plotted. **G)** Model of oRP preservation during stress in chaperone-stirred condensates.

To determine whether preserving oRPs during stress might promote fitness, we performed growth measurements of cells during recovery from heat shock (Figure 6E). We either left Sis1 in the nucleus or transiently depleted it and heat shocked the cells. After 20 minutes of heat shock, we allowed Sis1 to return to the nucleus in the cells in which it had been depleted and measured growth over time under non-heat shock conditions. In the Sis1+ cells, we observed immediate resumption of growth at a constant doubling rate that persisted until the cells plateaued after 300 minutes (Figure 6F). The cells in which Sis1 was transiently depleted showed slower growth and did not plateau until after 600 minutes (Figure 6F). Cells that were mock heat shocked showed no difference in growth in the Sis1+ and Sis1-depleted conditions (Figure S6C). This suggests that the reversibility of oRP condensates enabled by Sis1 contributes to resumption of rapid growth upon recovery from stress (Figure 6G).

## DISCUSSION

Upon exposure to a broad range of environmental stressors, cells rapidly inactivate ribosome biogenesis by repressing transcription of rRNA and RP mRNAs (Gasch et al., 2000; Gasch and Werner-Washburne, 2002; Sawarkar, 2022; Shore et al., 2021). Limiting production of ribosomes limits proliferation (Warner, 1999), providing a general mechanism for cells to simultaneously slow growth during stress and free up resources to mount an adaptive response. However, this rapid transcriptional repression is imperfectly coordinated with translational repression, potentially leaving newly synthesized RPs orphaned from rRNA during stress. Here we show that heat shock triggers the accumulation of oRPs at the periphery of the nucleolus in yeast and human cells. Rather than being degraded or forming toxic aggregates, oRPs form dynamic condensates which preserve the RPs for future use upon recovery from stress. oRP condensates retain their liquid-like dynamics in lysate and their reversibility in cells by virtue of the Sis1 and Hsp70 chaperones. We propose that a “chaperone-stirring” mechanism – in which Sis/Hsp70 activity prevents solidification of oRP condensates – enables cells to rapidly restart ribosome biogenesis and efficiently resume proliferation when conditions improve.

Like stress-induced condensates formed by the translation factors Pab1 and Ded1 (Iserman et al., 2020; Riback et al., 2017), oRP condensates appear to play an adaptive role in the yeast stress response. For Pab1 and Ded1, the ability to undergo temperature-dependent phase separation has been linked to fitness. In the case of oRPs, potential benefits of condensate formation include protection from degradation – thereby conserving the resources invested in synthesizing the RPs – and prevention of toxic gain of function effects. Moreover, the reversibility of oRP condensates may additionally contribute to fitness by enabling cells to rapidly resume ribosome biogenesis and proliferation.

Our data suggest that active engagement of Sis1 and Hsp70 with the condensates maintains the RPs in a usable state. We envision that a sudden buildup of oRPs drives initial condensation, but such assemblies would quickly become amorphous aggregates without Sis1 and Hsp70. Sis1 and Hsp70 in collaboration with other chaperones have been shown to efficiently disperse Pab1 condensates (Yoo et al., 2022), and Hsp70 has been implicated in repressing transcriptional condensates formed by the HSR regulator Hsf1 (Chowdhary et al., 2022). The additional role we identify here in maintenance of oRP condensates suggests that Sis1 and Hsp70 may broadly surveil and remodel biomolecular condensates during stress.

Basic questions about the composition, properties, and regulation of oRP condensates remain open. For example, the relationship between oRP condensates and the nucleolus – which has itself been interpreted as functioning as a proteostasis compartment (Frottin et al., 2019) – will be interesting to resolve in both the yeast and human cell contexts. With respect to the condensates themselves, we do not know the mechanisms that initiate their formation or trigger dispersal, we have not yet captured oRP condensates during the liquid-to solid-like transition, and we have not established a biophysical definition of the “chaperone stirring” mechanism. It will be particularly interesting to understand whether Hsp70 is acting as a multiple-turnover enzyme that repeatedly disrupts interactions among RPs or more as a holdase that blocks exposure of aggregation-prone regions on RPs.

Despite these unknowns, our data suggest a phenomenological model in which heat shock inhibits production of rRNA, resulting in the buildup and condensation of oRPs on the surface of the nucleolus (Figure 6G). Sis1 and Hsp70 actively maintain the oRPs in a usable state in the condensates until rRNA production resumes and the RPs are incorporated into nascent ribosomes. Supporting this model, cells are known to repress rRNA production during stress (Sawarkar, 2022; Shore et al., 2021), and inhibition of rRNA production has been shown to lead to accumulation of oRPs and activation of the HSR in the absence of environmental stress (Albert et al., 2019; Tye et al., 2019). While the mechanism by which heat shock inhibits transcription is unclear, it is known to be independent of the HSR (Rawat et al., 2021). Given that many nucleolar factors form reversible aggregates during heat shock (Wallace et al., 2015), perhaps heat shock triggers the condensation of rRNA biogenesis factors to inactivate rRNA production. These putative condensates of rRNA factors would trigger the downstream formation of oRP condensates. Such a “condensate cascade” would then culminate in the formation of additional condensates composed of the transcription factor Hsf1 and the transcriptional machinery that drive HSR activation (Chowdhary et al., 2022; Kainth et al., 2021). The chaperone machinery induced by the HSR could then disperse all these condensates to deactivate the HSR and resume growth in a concerted manner.

Our data suggest a direct connection between the biophysical properties of oRP condensates and fitness. We established this connection by depleting Sis1 from the nucleus, which solidified oRP condensates in lysate, rendered them irreversible in cells, and slowed proliferation upon recovery from heat shock. Loss of liquid-like properties has been associated with neurodegenerative disease-associated condensates (Alberti and Dormann, 2019; Alberti and Hyman, 2021), so it is possible that solidification of oRP condensates may play an analogous role in ribosomopathies. Alternatively, solid oRP aggregates may be nontoxic but simply unusable, interfering with the ability of chaperones to penetrate and facilitate rapid dispersal. In healthy cells, oRP condensates define proteostasis hubs that play an adaptive role in the stress response. The chaperone-stirring mechanism that we propose to sustain the liquid-like properties of the oRP condensates could function more broadly to transform potentially deleterious aggregates into benign condensates with adaptive potential.

## ACKNOWLEDGEMENTS

We thank V. Bindokas and C. Labno at the University of Chicago Integrated Light Microscopy Core (RRID: SCR_019197) for imaging assistance. We are especially grateful to B. Glick and lab members for the yeast HaloTag construct and advice on using JF dyes in yeast. We thank S. Kron for use of gel imaging instruments, and K. Lin for assistance aligning the Squires Lab custom wide-field setup and camera calibration. We also thank members of the Pincus, Squires, and Drummond labs for helpful discussions. J.A.M.B. acknowledges fellowship support from the Helen Hay Whitney Foundation. This work was supported by NIH grants R01 GM138689 to D.P., R35 GM144278 to D.A.D., support from the Neubauer Family Foundation to A.H.S., and NSF QLCI QuBBE grant OMA-2121044 to D.P. and A.H.S.

## AUTHOR CONTRIBUTIONS

Conceptualization: A.A. and D.P.; Methodology: A.A., R.G., O.C.S., J.A.M.B., K.H., S.K., A.H.S.; Formal Analysis: A.A., O.C.S., K.H., A.H.S., D.P.; Investigation: A.A., R.G., O.C.S., J.A.M.B., S.K., K.D., S.L-W.; Resources: A.A., J.A.M.B., D.A.D., A.H.S., D.P.; Writing – original draft: A.A., D.P.; Writing – Review and Editing: all authors.; Visualization: A.A., O.C.S., K.H., D.P.; Supervision: D.A.D., A.H.S., D.P.; Funding Acquisition: D.A.D., A.H.S., D.P.

## METHODS

### Yeast strain construction and cell growth

Yeast strains used in this study are catalogued in Table S1. All strains are derivatives of W303, and fluorescent protein and epitope tags are integrated into the genome using scarless CRISPR/Cas9 gene editing unless otherwise stated. Yeast were cultured in rich glucose medium (YPD) or appropriate synthetic complete (SC) drop-out media at 30°C. For lattice light sheet imaging, cells were cultured in SDC-riboflavin and folic acid to minimize autofluorescence.

### Pulse labelling of newly synthesized or mature Halo-tagged ribosomal proteins (yeast)

To visualize newly synthesized ribosomal proteins, corresponding strains with ribosomal proteins fused with HaloTag7 (codon optimized for S. Cerevisiae) were grown to O.D 0.3-0.6 and incubated with non-fluorescent Halo tag ligand 7-bromo-1-heptanol (7BRO, 100μM, VWR #AAH54762) for 10 min to irreversibly mask all the pre-existing ribosomal proteins. Samples were washed three times with fresh media (5X volume), shaking for 1 min between each wash. Cells were incubated with the Janelia Fluor 646 Halo tag ligand (JF646, 1μM, Promega #GA1120) in media at 30°C or 39°C for 7 min followed by washing (5X volume) with prewarmed media (30°C/39°C) for additional 3 min prior to imaging. To label and visualize mature ribosomal proteins, yeast were incubated with 10 μM JF646 Halo dye for 10 mins. Excess dyes was washed away, and further labelling with dye is prevented by addition of 100 μM 7BRO.

### Acute depletion of Ifh1 and RNA sequencing

Genomic *IFH1* was fused in frame with mini auxin inducible degron (miniAID) at its C termini using CRISPR/Cas9 gene editing. Estradiol inducible OsTIR1(F74G) was introduced at the *leu2* locus. Acute and non-leaky depletion of Ifh1 is achieved by co-treatment with β-estradiol (1μM, Sigma #E8875) and 5ph-IAA (5μM, MedChemExpress #HY-134653) for 30 min. Sequencing of polyA+ mRNA from yeast was performed at the Northwestern University Genomics Core and analyzed described (Feder et al., 2021; Zheng et al., 2016).

### Transient Sis1 depletion and quantitative growth assay

1 μM rapamycin (Sigma #R8781) was used to anchor away Sis1-FRB in a rapamycin-resistant background (*TOR1-1 fpr1Δ*) with Rpl13a-FKBP (Feder et al., 2021). Three biological replicates of the Sis1-FRB/Rpl25-FKBP strain were grown at 30°C overnight in 2xSDC media, then diluted to OD 0.05 in the morning. Cells were growth for 5 hours at 30°C until thee OD reached 0.2. Each biological replicate was split into two tubes, corresponding to DMSO pretreated followed by HS (39°C, 20 min) or rapamycin pre-treated (1uM, 10 min) followed by HS (39°C, 20 min). All samples were then washed three times with 2xSDC media, resuspended in 2xSDC to final OD 0.1, and OD measurements taken an automated plate reader every 20 minutes for the next 24 hours while shaking at 30°C.

### Human cell culture

HCT116 and HCT116 RPL26-Halo were grown in Dulbecco’s modified Eagle’s medium (DMEM, high glucose and pyruvate) supplemented with 10% (vol/vol) fetal bovine serum, 1X GlutaMAX and 100 units of penicillin and streptomycin, and maintained in a 5% CO_2_ incubator at 37 °C. All cell lines were found to be free of mycoplasma using the MycoAlert^®^ PLUS Mycoplasma Detection Kit (Lonza #LT07-703).

### Conditioned media heat shock treatment

Heat shock with conditioned media was performed essentially as described (Mahat and Lis, 2017). Briefly, two sets of cell plates were grown identically to 70-80% confluence. Conditioned media from the first cell plate was collected and prewarmed to 43°C in bead bath. The cells on this plate were discarded. The conditioned media was heated to 43°C, while media from the second cell plate were discarded and replaced with heated conditioned media collected from the first cell plate and immediately placed in a 43°C incubator with 5% CO_2_.

### Pulse labelling of new or mature Halo-tagged ribosomal proteins (human cell lines)

HCT116-RPL26-Halo strain were seeded on 24-well plates 30 hours prior to the experiment. To visualize newly synthesized RPL26 cells, pre-existing RPL26 were masked by incubation with non-fluorescent irreversible Halo ligand (7BRO, 100 μM) for 15 min. Cells were washed with complete DMEM for 3 times under shaking for 2 min. Cells were then treated with JF646 Halo dye (1 μM) mixed with prewarmed conditioned media and heat shocked as indicated. Conversely to label pre-existing RPL26, cells were incubated with JF646 Halo dye (10 μM) for 15 min. After extensive washing, further labelling was prevented by addition of 100 μM 7BRO.

### Immunofluorescence of HCT116 cells

HCT116 RPL26-Halo cells were seeded at 0.8 × 10^5^ cells/mL on poly-lysine-coated coverslips 30 hour prior to the experiment. Cells were fixed with 4% paraformaldehyde, 4% sucrose in 1XPBS for 5 min. Fixed cells were washed with 1XPBS and fixation was quenched with 125 mM glycine. Thereafter, cells were permeabilized with 0.1% Triton-X100 followed by blocking with 5% normal goat serum in 1X PBS. DnaJB6 was stained by incubating cells overnight with 1:100 dilution polyclonal rabbit DnaJB6 antibody (ThermoFisher #PA5-27577). Nucleolus was stained by incubating with 1:1000 dilution of mouse monoclonal NPM1 antibody (ThermoFisher #FC-61991). Cells were then washed with 1XPBS and fluorescently labeled with goat–anti-rabbit 488 and goat–anti-mouse 594 secondary antibodies (Invitrogen). Cells were then washed with 1XPBS, mounted on microscope slides using ProLong Gold Antifade Mountant with DAPI (Invitrogen #P36930) and imaged the following day. Images were taken using a 3i Marianas Spinning Disk Confocal with 100X oil objective and pseudo colored with Fiji.

### Lattice light-sheet imaging and analysis

Lattice light-sheet imaging of live yeast was executed using a phase 2 system manufactured by Intelligent Imaging Innovations (3i) and run with SlideBook 6.0 software. The design is a commercially produced clone of the original (Chen et al., 2014) with greater automation and stability. The imaging camera used was a Hamamatsu Fusion chilled sCMOS with annulus mask set for a 20 μm beam length (outer NA, 0.55; inner NA, 0.493) with 400-nm thickness, with dither set at 9 μm. Temperature was controlled by a built-in Peltier device (empirically set using a probe thermometer to indicated temperatures). Optics were aligned daily before the experiment, and the bead PSFs (at the respective temperature and media) were collected before imaging to ensure that the setup has consistent resolution in XYZ, in addition to providing the PSF in each channel for deconvolution. Graphics processing unit–based Richardson-Lucy deconvolution was used to denoise and enhance z stack images using measured PSFs or theoretical PSF via Brian Northan’s “Ops” implementation (https://github.com/imagej/ops-experiments). 3D reconstructions and videos were assembled using MPI-CBG developed ClearVolume plugin of Fiji (Royer et al., 2015; Schindelin et al., 2012).

### APEX2 proximity labeling

Sis1-Apex2 strains were grown in 300 ml SDC media to OD 0.3-0.6. Cells were heat shocked at 39°C for 15 mins by immediate mixing with 300ml of pre-warmed media at 50°C. Post-treatment cells were fixed with 1% Paraformaldehyde for 5 min. The fixation was quenched with 125mM glycine and cells were vacuum filter collected and washed with potassium phosphate buffer saline (KpiS) buffer three times. Cells were resuspended in KpiS Buffer supplemented with 100T zymolyse to lyse cell wall. Cell membranes were permeabilized with 0.1% TritonX100 in KpiS Buffer. Spheroplast were incubated with biotin phenol (0.5 mM) for 20 mins at 30°C. To activate APEX2 labelling reactions, Hydrogen peroxide (1 mM) was added to the medium for 1 min. For background correction, one set without peroxide treatment was identically processed. The reactions were immediately quenched with a cocktail of Trolox (2.5 mM), Sodium Ascorbate (10 mM) and Sodium azide (10 mM) in 1X PBS. Cells were harvested by centrifugation and the pellet was flash frozen in liquid nitrogen. Cell pellets were cryo-milled, and the lysate was dissolved in 1X RIPA buffer. Proteins were precipitated with ice cold methanol at -80°C for overnight. Protein precipitants were dissolved in 2% SDS RIPA Buffer solution under 95°C for 20 mins. Biotinylated proteins were affinity captured with streptavidin magnetic beads. Washed with RIPA ×2, 1M KCl, 0.1M Na2CO3, 2M urea in HEPES buffer ×2 and RIPA ×2. Completely dried beads were then resuspended in 50 μl of hexafluoro isopropanol (HFIP), incubated for 5 min with shaking and eluted by spinning. Fifty microliters of HFIP were added to the beads again, and the eluates were combined and dried in a speedvac for 10 min. This sample was used for LC/MS analysis.

### Sis1 co-IP from cells labeled for mature and new ribosomal proteins

Corresponding Rp-Halo strains with Sis1-3xFlag were grown to mid log phase in 300 ml SDC media. Cells were labelled for either mature or new ribosomal proteins as described above (pulse labeling section). Cells were kept unstressed, or heat shocked (39°C) for the indicated time by mixing with equal volume media at 50°C. Following treatment, cells were harvested by vacuum filtration and flash frozen in liquid nitrogen. Cells were lysed by cryo-milling at 6×90sx30Hz pulses in a Retsch MM100 mixer mill under liquid nitrogen. Samples were dissolved in physiological buffer (20mM NaCl, 50mM KCl, 150mM K+Glutamate, 50mM HK_2_PO_4_, 0.5mM MgCl_2_, 25mM HEPES-KOH, 25mM MES pH 7.4). Sis1-3xFlag were immunoprecipitated using anti-Flag M2 Magnetic beads (Sigma #M8823) from the cell lysates. Bead-bound complexes were released with competition for 3xFlag peptide (Sigma #F4799). IP eluates were run on SDS-PAGE gel and visualized using a fluorescence gel imager. Gels were then blotted and probed with anti-Flag (Sis1). Co-IP experiments were performed in triplicate.

### Proteomics sample preparation

Samples were run on SDS-PAGE and gel bands were subjected for in-gel digestion. Gel bands were washed in 100mM Ammonium Bicarbonate (AmBic)/Acetonitrile (ACN) and reduced with 10mM dithiothreitol at 50°C for 45 min. Cysteines were alkylated with 100mM iodoacetamide in the dark for 45 min in room temperature (RT). Gel bands were washed in 100mM AmBic/ACN prior to adding 1μg trypsin (Promega #V5111) for overnight incubation at 37°C. Supernatant containing peptides were collected into a new tube. Gel pieces were washed with gentle shaking in 50% ACN/1% FA at RT for ten minutes, and supernatant was collected in the previous tubes. Final peptide extraction step was done with 80% ACN/1% FA, and 100% ACN, and all supernatant was collected. The peptides were dried at speedvac and reconstituted with 5% ACN/0.1% FA in water before injecting into the LC-MS/MS.

### LC-MS/MS analysis

LC-MS/MS analysis was performed at the Nothwestern University Proteomics Core. Peptides were analyzed by LC-MS/MS using a Dionex UltiMate 3000 Rapid Separation nanoLC coupled to a Q-Exactive HF (QE) Quadrupole-Obitrap mass spectrometer (Thermo Fisher Scientific Inc, San Jose, CA). Samples were loaded onto the trap column, which was 150 μm x 3 cm in-house packed with 3 um ReproSil-Pur® beads. The analytical column was a 75 um x 10.5 cm PicoChip column packed with 3 um ReproSil-Pur® beads (New Objective, Inc. Woburn, MA). The flow rate was kept at 300nL/min. All fractions were eluted from the analytical column at a flow rate of 300 nL/min using an initial gradient elution of 5% B from 0 to 5 min, transitioned to 40% over 100 min, 60% for 4 mins, ramping up to 90% B for 3 min, holding 90% B for 3 min, followed by re-equilibration of 5% B at 10 min with a total run time of 120 min. Mass spectra (MS) and tandem mass spectra (MS/MS) were recorded in positive-ion and high-sensitivity mode with a resolution of ∼60,000 full-width half-maximum. The 15 most abundant precursor ions in each MS1 scan were selected for fragmentation by collision-induced dissociation (CID) at 35% normalized collision energy in the ion trap. Previously selected ions were dynamically excluded from re-selection for 60 s. Proteins were identified from the MS raw files using the Mascot search engine (Matrix Science, London, UK. version 2.5.1). MS/MS spectra were searched against the SwissProt Saccharomyces cerevisiae database. All searches included carbamidomethyl cysteine as a fixed modification and oxidized Methionine, deamidated asparagine and aspartic Acid, and acetylated N-term as variable modifications. Three missed tryptic cleavages were allowed. A 1% false discovery rate cutoff was applied at the peptide level. Only proteins with a minimum of two peptides above the cutoff were considered for further study. Identified peptides/protein were visualized by Scaffold software (version 5.0, Proteome Software Inc., Portland, OR).

### Polysome profiling

Polysome analyses by sucrose gradient fractionation were performed as described (Aboulhouda et al., 2017). Rpl26a-Halo strains were grown in SDC media (160 ml) to mid-log phase (OD 600 ∼0.5). Pre-existing ribosomes were masked by incubation with 7BRO (100 μM, 10 mins). Cells were washed with fresh media (5x volume) 3 times under shaking for 2 min. Newly synthesized Rpl26a were labelled by incubation with JF646 Halo ligand (2 μM). Following which cells were heat shocked at 39°C (10 min) by mixing with equal volume media at 50°C. One half of the cells were harvested by vacuum filter and denoted as heat shock sample. Another half were allowed to recover at 30°C in the presence of blocker (100 μM) for 30 min. Similarly, this sample was vacuum filter collected and marked as recovery sample.

The cell pellet was transferred to a 2 ml Eppendorf “Safe-Lok” tube and flash frozen in liquid nitrogen. Cells were lysed by a pre-chilled 7 mM stainless steel ball (Retsch #05.368.0035) with 4×90sx30Hz pulses in a Retsch MM100 mixer mill, chilling in liquid nitrogen (LN) between pulses. Sample was resuspended in 500 μl lysis buffer (20mM HEPES-KOH (pH 7.4), 100mM KCl, 5mM MgCl2, 200μg/mL heparin (Sigma #H3149), 1% triton X-100, 0.5mM TCEP, 100μg/mL cycloheximide, 20 U/ml superase-IN (Invitrogen #AM2696), 1:200 Millipore protease inhibitor IV #539136). The lysate was clarified by centrifugation at 3000 g for 30 s, and the supernatant was transferred to new tube and aliquots were flash frozen in LN.

A 10–50% continuous sucrose gradient in polysome gradient buffer (5mM HEPES-KOH (pH 7.4), 140mM KCl, 5mM MgCl2, 100μg/ml cycloheximide, 10 U/ml superase-in, 0.5 mM TCEP) was prepared in SW 28.1 tubes (Seton #7042) using a Biocomp Gradient Master and allowed to cool to 4°C. 200 μl of clarified lysate was loaded on top of the gradient, and gradients were spun in a SW28.1 rotor at 27500 rpm for 3.5 hr at 4°C. Gradients were fractionated into 15 fractions using a Biocomp Piston Gradient Fractionator with UV monitoring at 254 nm, and fractions were flash frozen in LN. UV traces were normalized to the total signal starting with the 40S peak. For in gel-fluorescence samples were treated with 0.02% sodium deoxy cholate and precipitated by adding 10% TCA for 1 hr on ice. Pellets were extensively washed with ice-cold acetone, and then resuspended in 2X Laemmli sample buffer.

### Cell-free lysate droplet assay

Corresponding newly synthesized or matured ribosomal protein-stained yeast cultures were heat shocked, filter collected, cryo-milled and dissolved in 1:2 (w/v) in physiological buffer (20mM NaCl, 50mM KCl, 150mM K+Glutamate, 50mM HK_2_PO_4_, 0.5mM MgCl_2_, 25mM HEPES-KOH, 25mM MES pH 7.4, 5mM PMSF and 1:200 Millipore Protease Inhibitor Cocktail). Cellular debris and un-lysed cells were clarified by centrifugation at 6000 rpm for 5 min. Cleared lysates were immediately loaded onto a homemade microfluidic chamber comprising a glass slide or coverslip with a top coverslip attached by two parallel strips of double-sided tape to form a single central channel (Roy et al., 2008). Slides were then imaged with a custom wide-field microscope with high NA objective (Olympus PlanApo N 60X/1.42), excited with a 637nm continuous laser at ∼10 W/cm2 (Coherent Obis LX) and sCMOS camera detection (Photometrics Prime95B) at 66 fps. Imaged condensates have settled on the glass coverslip.

### Sis1 nucleolar proximity analysis

From the maximum intensity projections, nuclei were segmented by drawing oval masks around peri-nuclear ring of Sec61-Halo using Fiji software. This was used as an input to custom-written Python code, which carried out the rest of the analysis. For each nucleus, the geometric center was computed from the mask, and a family of lines drawn at varying angles to bisect the nuclear area into two halves. For each nucleus, the fold-change in Nsr1 and Sis1 intensity between the two halves was computed, and the angle with a maximal fold-change in Nsr1 intensity was identified as splitting the nucleus into Nsr1-proximal and Nsr1-distal. The fold-change in Sis1 signal along the bisecting angle was recorded for each nucleus.

### Colocalization analysis

To quantify colocalization between Sis1 and Rps signal, Coloc2 plugin of Fiji was utilized. Minimum error thresholding (MET) method coupled with watershed algorithm were used to generate mask for individual cell. Automatic Otsu (maximum between-cluster variance) thresholding was used for pixel intensity-based mask creation and segmentation of high intensity peri nucleolar Sis1 signal and cytosolic foci from the low intensity diffused signal. Mander’s overlap coefficient was calculated to determine the fraction of the mature or new form of RP signals overlap with Sis1 signals.

For human cells, colocalization analysis of DnaJB6 with RP signals with the Coloc2 plugin of FIJI was also utilized. Huang threshold method coupled with fill hole and watershed algorithm was utilized upon DAPI signal to create a mask and segment nuclei. Created masks were binary dilated to segment the cytosol. The ROI generated was used to calculate the Mander’s overlap coefficient between RPL26 (mature or new) and DnaJB6 signal.

### Yeast cytosol image segmentation

Deconvolved z stack images were maximum intensity projected. Estimated mask for whole cell were obtained using the Minimum error thresholding (MET) methods coupled with watershed algorithm upon Sis1 signal. Nsr1 signal were used for segmentation of yeast nucleoli. Nucleolar mask was binary dilated to enlarge and cover the nucleolar proximity region. Cytosolic signals were extracted by subtracting the nucleoalar+perinucleolar signal (approx. nuclei) with whole cell signal. Fraction cytosolic is determined by dividing the integrated density of whole cell over cytosol signal.

### Human cell cytosol image segmentation

Z series images were background subtracted and projected over 2D for maximum intensity. DAPI signal was used to mask and segment the nuclei using Li threshold coupled with watershed and fill hole algorithm (Fiji). DAPI mask were binary dilated and segment for whole cell were created. Fraction cytosolic RPL26 signal was calculated by dividing the whole cell integrated density with cytosol intergraded density.

### Condensate pixel variance analysis

To determine the relative dynamicity of the condensates using videos taken with a wide-field microscope, the time-dependent variance in condensate brightness was quantified. In comparison to the solid-state condensates, the liquid-like condensates were found to exhibit higher Poisson-normalized variance in pixel brightness over time, consistent with higher mobility of heterogeneously labeled contents, such as aggregates of fluorescent protein, moving around within a liquid-like environment. The time-dependent depletion of ATP or tethering of heat shock proteins was found to produce a normalized variance more consistent with a solid-like state, indicating that ATPases such as Hsp70 might be involved in maintaining a liquid-like state. Analysis was undertaken for videos of condensates under identical illumination using a bespoke Python script.

Pixel count values were converted to photon counts according to a previously calibrated gain matrix unique to the sCMOS camera used (Photometrics Prime95B). Converted videos were cropped to include only areas within the condensate. For each pixel at location (*i, j*) in the condensate, normalized pixel variance over time 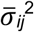 was calculated from the photon counts observed in each of *N* frames, *x*_*ij*_(*t*), and the expected variance of the signal according to a Poisson distribution of photon counts, 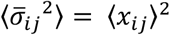, Equation 1.

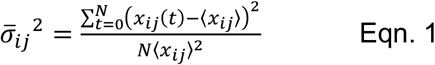

The resulting two-dimensional array of normalized variances for each condensate, ***σ***)^**2**^, was generated and displayed as a heatmap. Normalized variance values from multiple condensate videos of each condensate type were aggregated to enable robust comparison among treatment conditions, displayed as a violin plot. The expected normalized variance for solid objects is 1, since any variability in the signal would be due to shot noise generated from photon counting by the detector. Similarly, fluctuations above the expected shot noise variance would have values greater than 1, indicating that the object is exhibiting fluctuations in addition to those predicted from shot noise.

---

**Figure S1.**
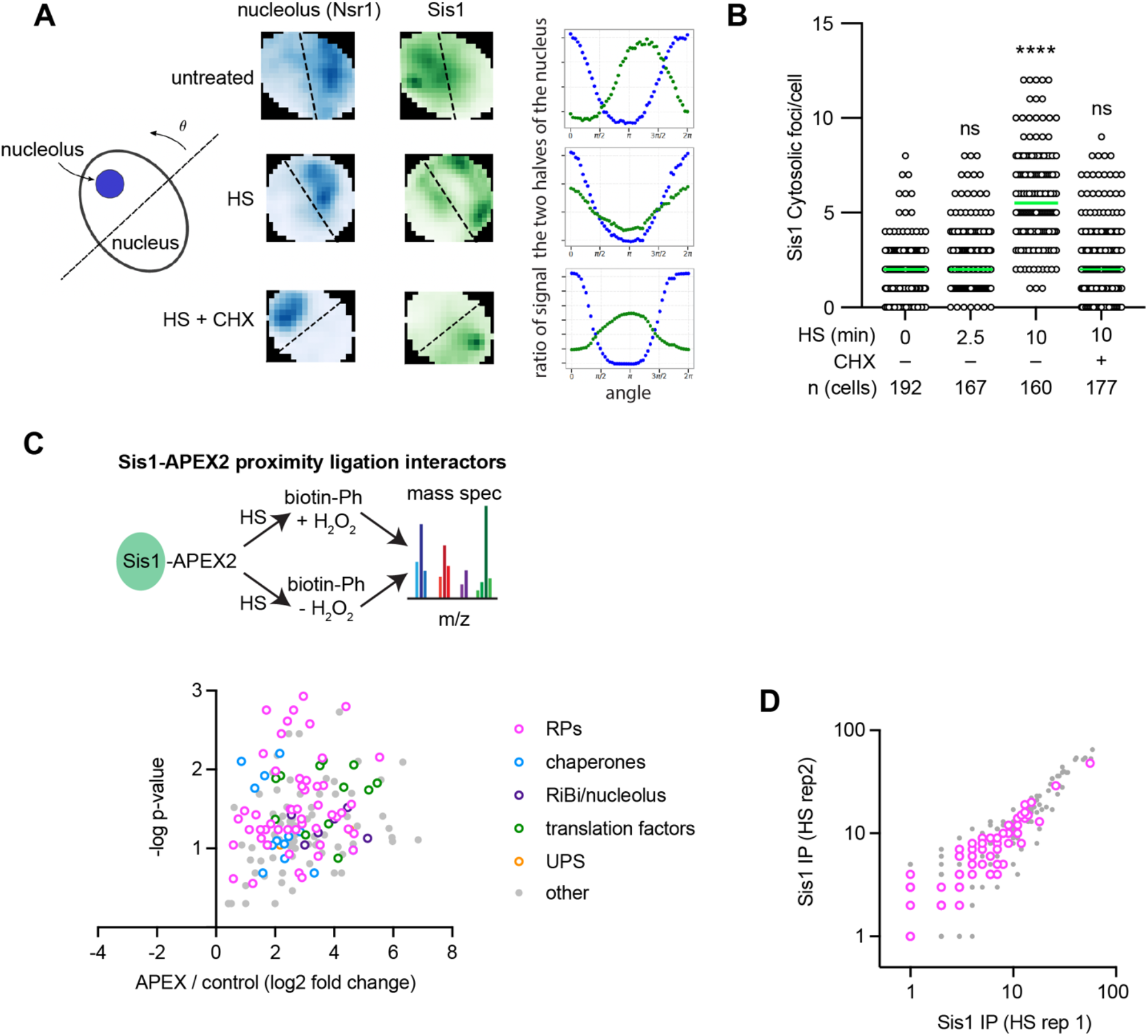
Sis1 localization and interactions during heat shock. **A)** Left: Schematic of how to bisect the nucleus with the nucleolus on one side by finding the line angle with the maximum difference in signal of the nucleolar marker in the two halves. Middle: Representative 2D projections of cells showing Nsr1 (blue) to mark the nucleolus and Sis1 (green). Line is set to maximize the difference in nucleolar signal and the ratio of Sis1 in the two halves is calculated. Right: Sis1 ratio as a function of the line angle rotated as depicted in the schematic to the left. **B)** Quantification of Sis1 cytosolic foci per cell in the conditions listed. Foci were identified using the FindFoci plugin in ImageJ. Statistical significance was determined by Brown-Forsythe and Welch ANOVA test with multiple comparisons. \*\*\*\**P* < 0.0001. **C)** Volcano plot of Sis1-APEX2 interactors during heat shock. **D)** Biological replicates of Sis1-3xFlag IP interactors.

**Figure S2.**
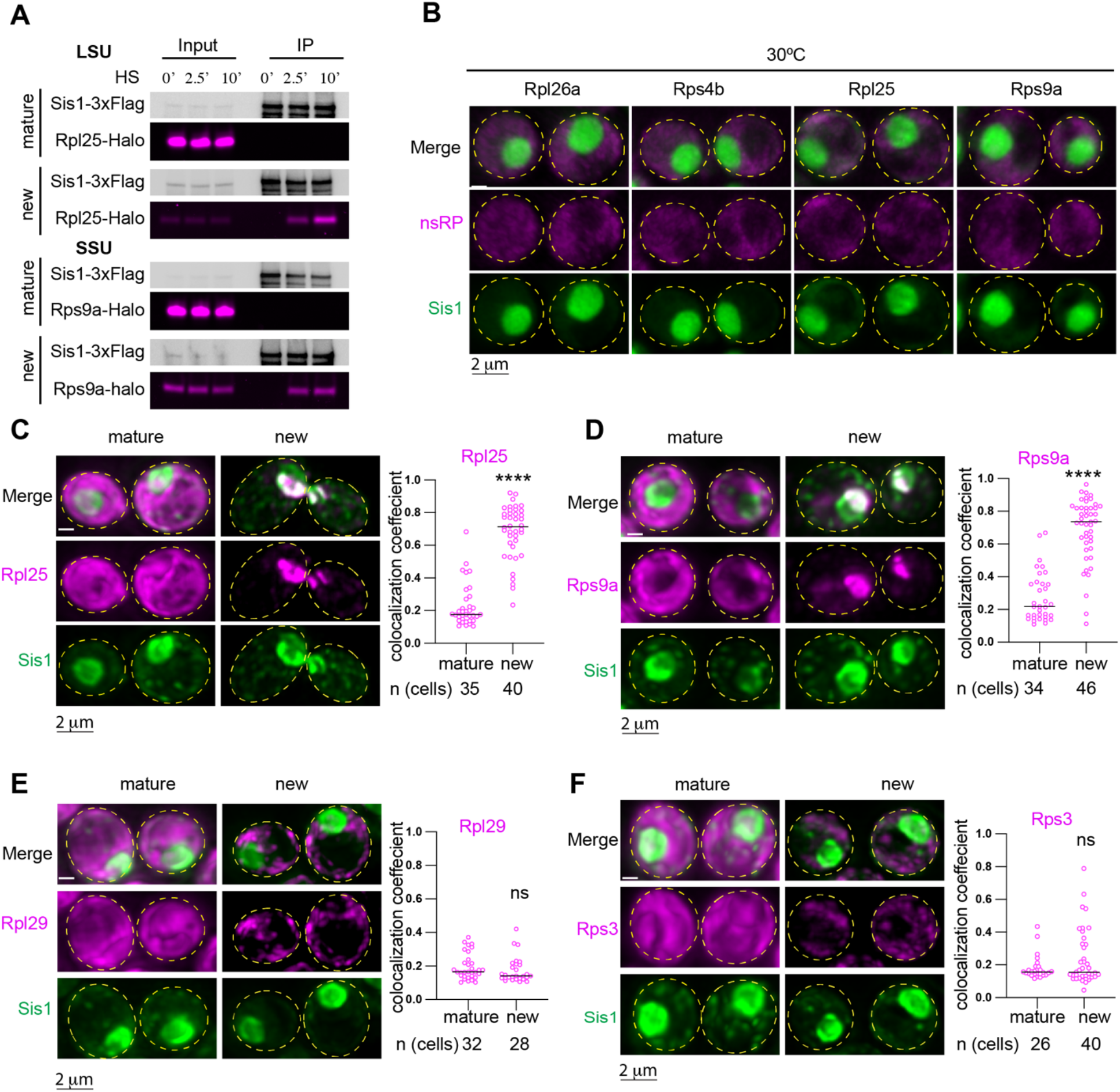
Interaction and localization of pulse-labeled ribosomal proteins with Sis1. **A)** IP of Sis1-3xFlag and either mature or new Rpl25-Halo and Rps9a-Halo from cells left unstressed or heat shocked at 39°C for the indicated times. **B)** In the absence of heat shock, pulse-labeled RPs localize immediately to the cytosol. **C)** Left Panel: Lattice light sheet live imaging of yeast under heat shock (39°C, 10 min) expressing Sis1-mVenus and labeled for either new or mature Rpl25-Halo. Right Panel: Dot plot representing the colocalization coefficient (Mander’s overlap coefficient) of Sis1-mVenus with either mature or new Rpl25-Halo in heat shocked cells (39°C, 10 min). **D)** As in (C) but for Rps9a-Halo. **E)** As in (C) but for the late joining subunit Rpl29-Halo. **F)** As in (C) but for the late joining subunit Rps3-Halo. *P* values were calculated with a two-tailed Welch’s t-test. \*\*\*\**P* < 0.0001. ns, (non-significant).

**Figure S3.**
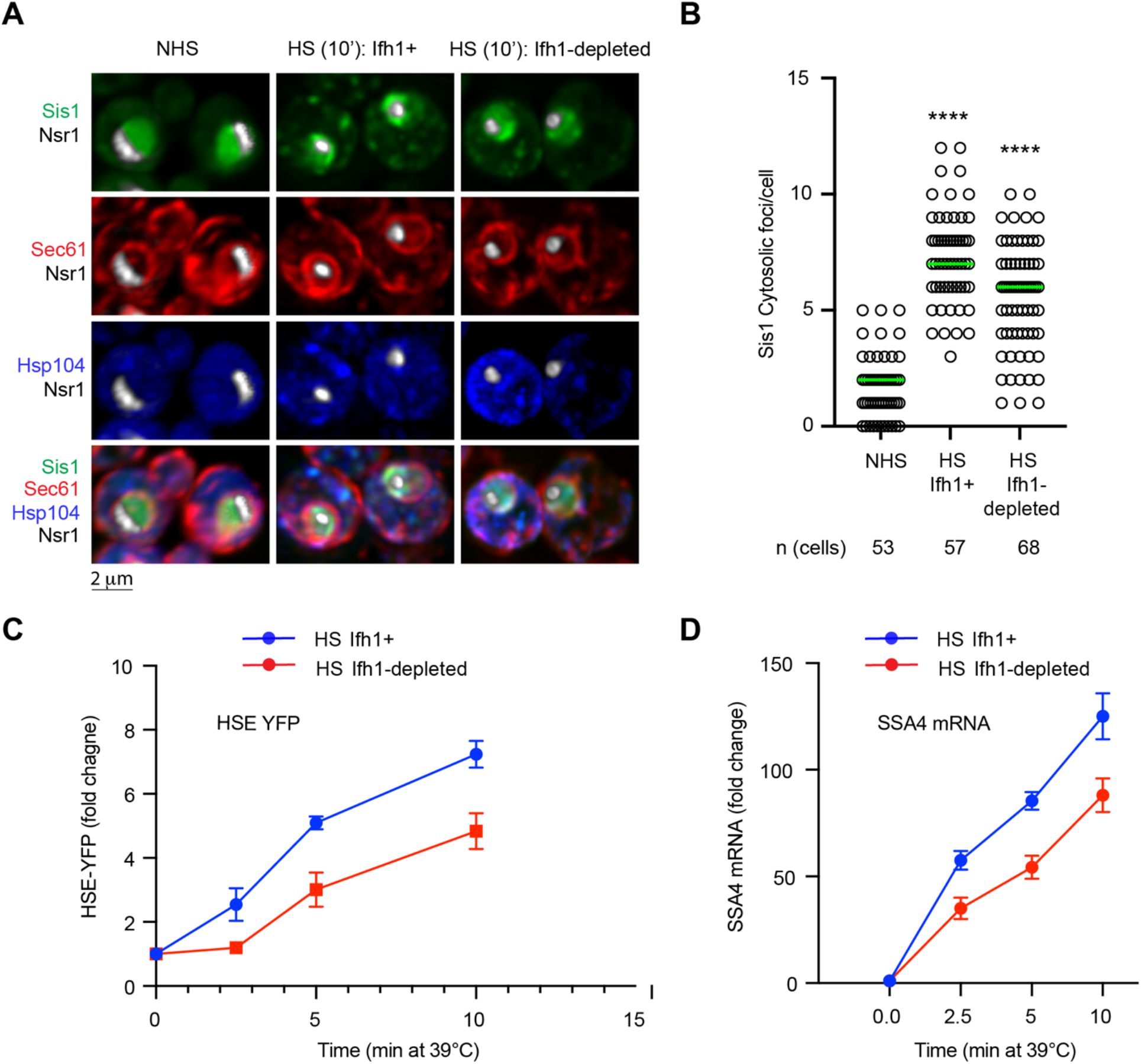
Cell biological and transcriptional effects of Ifh1 depletion. **A)** LLS live-imaging of yeast cells with endogenously tagged Sis1-mVenus (green), Hsp104-TFP (blue), Sec61-Halo (red) and Nsr1-mScarletI (white) under non-stress (30°C) and heat shock (39°C, 10 min) in the absence and presence of Ifh1 depletion. **B)** Quantification of Sis1 cytosolic foci per cell in the conditions shown in (A). Statistical significance was determined by Brown-Forsythe and Welch ANOVA test with multiple comparisons. \*\*\*\**P* < 0.0001. **C)** HSE-YFP reporter heat shock time course showing reduced HSR induction when Ifh1 is depleted. **D)** RT-qPCR of the HSR target gene transcript *SSA4* over a heat shock time course in the absence and presence of Ifh1 depletion.

**Figure S4.**
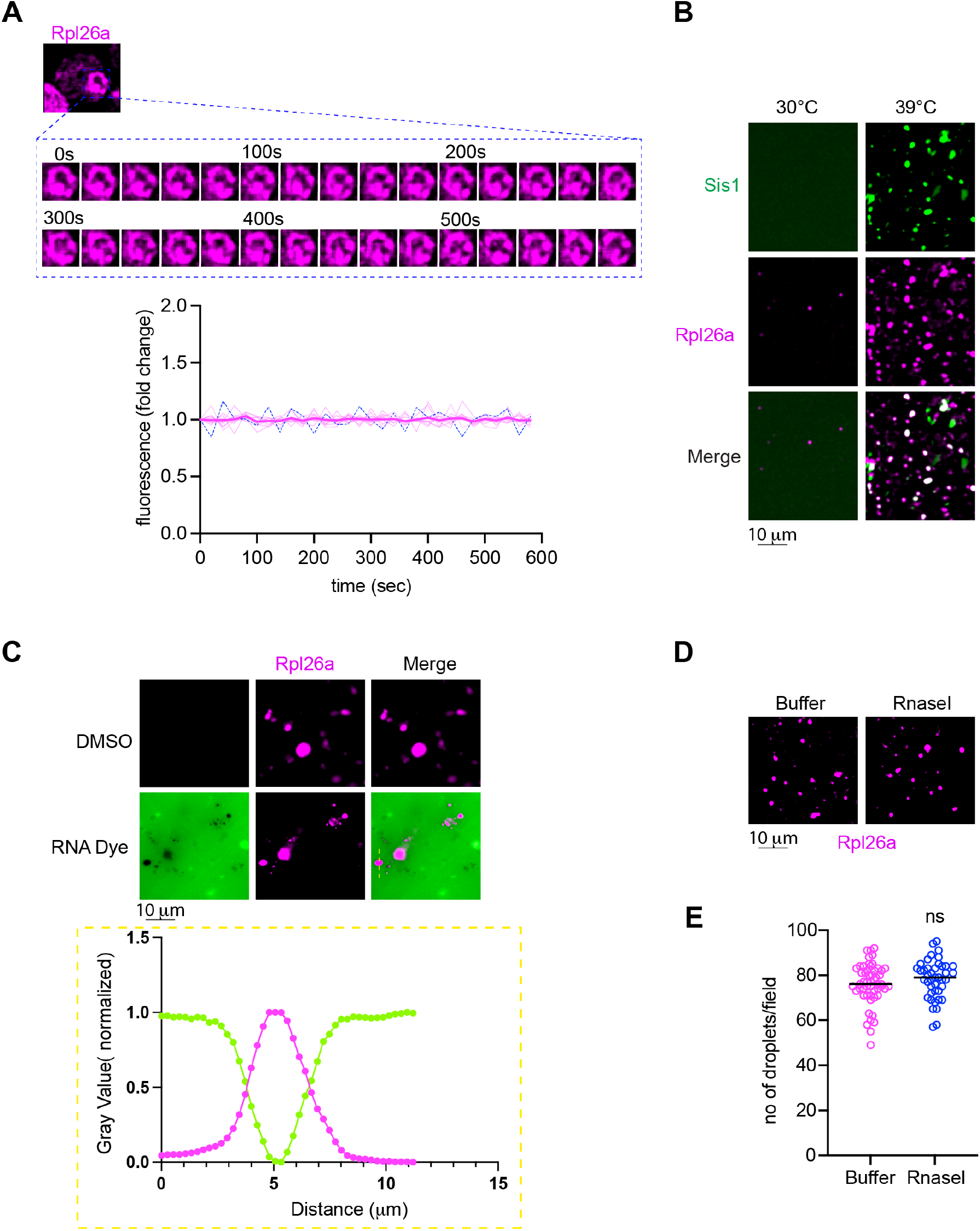
oRP condensates are stable, heat shock-dependent, and RNA-free. **A)** oRP proteins are stable (not degraded) in condensates in cells. **B)** oRP condensates are more abundant in lysate from heat-shocked cells. **C)** RNA dye (SYTO RNASelect, 0.5mM, 10 min) is excluded from the oRP condensate. **D)** oRP condensates are resistant to RNaseIf (5 units/μl, 15 min, RT). **E)** Quantification of number of droplets per field in buffer or RNaseIf treatment to the lysate. *P* values were calculated with a two-tailed Welch’s t-test. ns, (non-significant).

**Figure S5.**
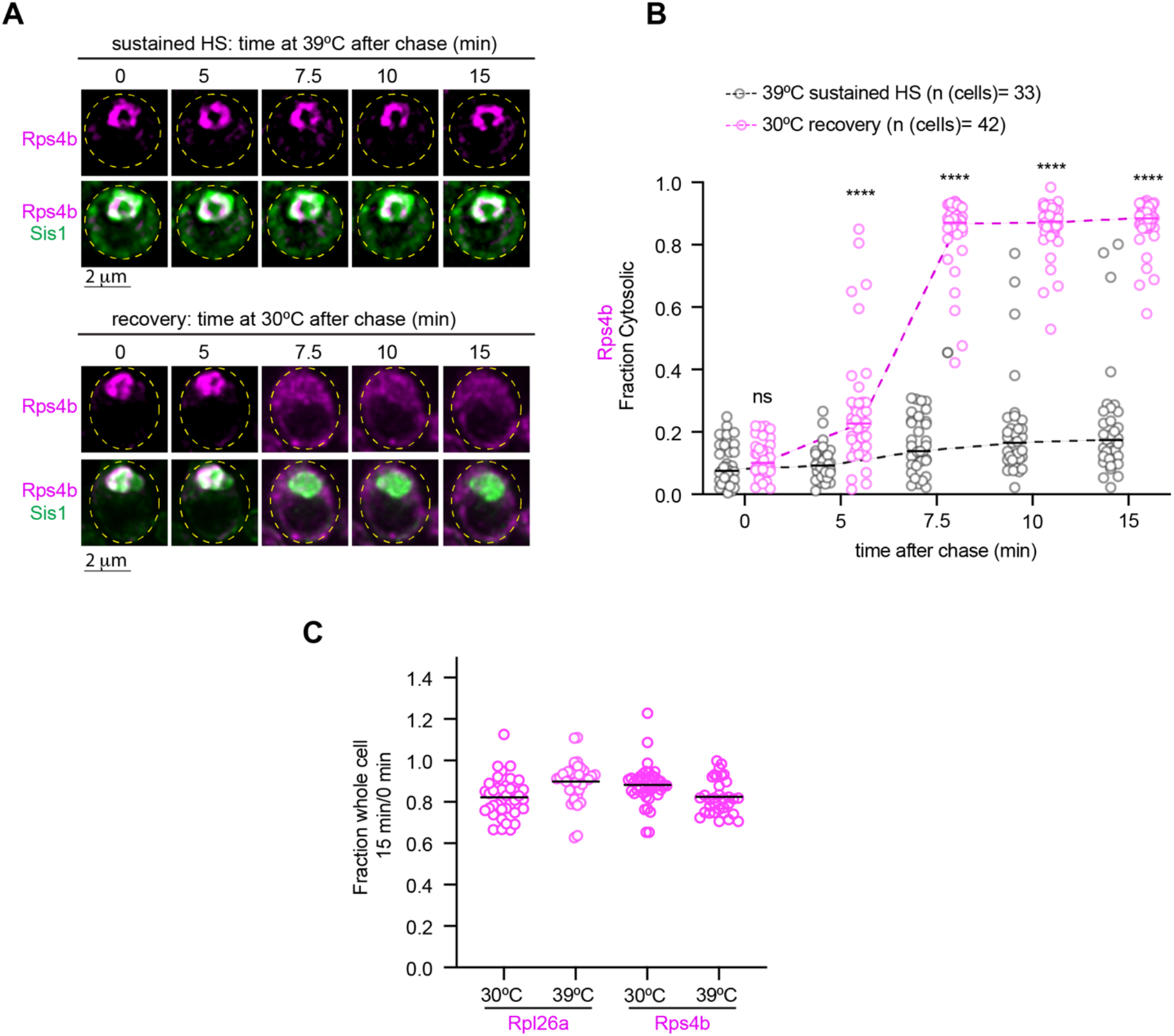
oRPs in condensates are not degraded and are transported to the cytosol upon recovery. **A)** Live cell time lapse imaging of the spatial distribution of oRps4b (magenta) and Sis1-mVenus (green) during sustained heat shock and recovery **B)** Quantification of the fraction of cytosolic oRps4b signal under sustained HS or recovery. Statistical significance was determined by Brown-Forsythe and Welch ANOVA test with multiple comparisons. \*\*\*\**P* < 0.0001, ns (non-significant). **C)** Fraction of total pulse labeled Rpl26a or Rps4b remaining after 15 minutes at each temperature

**Figure S6.**
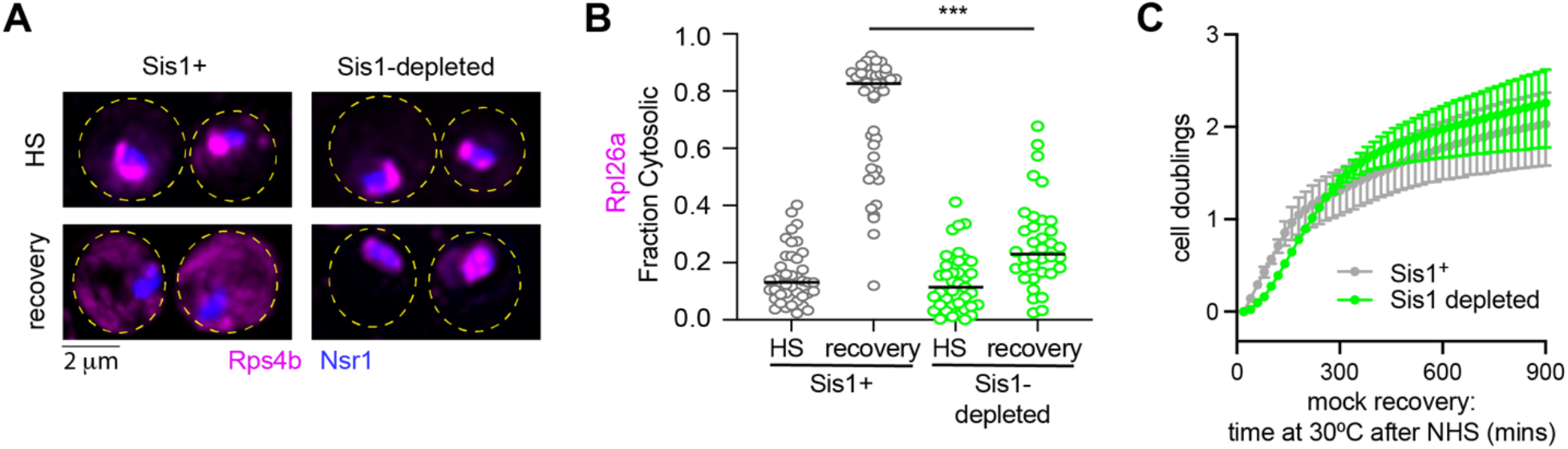
oRP condensate reversibility depends upon Sis1 availability. **A)** LLS live cell imaging of oRps4b (magenta) and the nucleolar marker Nsr1 (blue) during heat shock and recovery in the absence or presence of Sis1 depletion. **B)** Quantification of the fraction of cytosolic Rpl26a under sustained HS or recovery in the absence or presence of Sis1 depletion. *P* values were calculated with a two-tailed Welch’s t-test. \*\*\*\**P* < 0.0001. **C)** Growth curves of cells following “recovery” from mock heat shock (cells were not heat shocked, just grown at 30ºC, treated with 30ºC media and then grown at 30ºC) in the presence or absence of transient Sis1 depletion as depicted in Figure 6.

